# Mechanisms of cilia regeneration in *Xenopus* multiciliated epithelium *in vivo*

**DOI:** 10.1101/2023.06.14.544972

**Authors:** Venkatramanan G. Rao, Vignesharavind Subramanianbalachandar, Magdalena M. Magaj, Stefanie Redemann, Saurabh S. Kulkarni

## Abstract

Cilia regeneration is a physiological event, and while studied extensively in unicellular organisms, it remains poorly understood in vertebrates. In this study, using *Xenopus* multiciliated cells (MCCs) as a model, we demonstrate that, unlike unicellular organisms, deciliation removes the transition zone (TZ) and the ciliary axoneme. While MCCs immediately begin the regeneration of the ciliary axoneme, surprisingly, the assembly of TZ is delayed. However, ciliary tip proteins, Sentan and Clamp, localize to regenerating cilia without delay. Using cycloheximide (CHX) to block new protein synthesis, we show that the TZ protein B9d1 is not a component of the cilia precursor pool and requires new transcription/translation, providing insights into the delayed repair of TZ. Moreover, MCCs in CHX treatment assemble fewer (∼ 10 vs. ∼150 in controls) but near wild-type length (ranging between 60 to 90%) cilia by gradually concentrating ciliogenesis proteins like IFTs at a select few basal bodies. Using mathematical modeling, we show that cilia length compared to cilia number influences the force generated by MCCs more. In summary, our results question the requirement of TZ in motile cilia assembly and provide insights into how cells determine organelle size and number.

## INTRODUCTION

Cilia perform critical functions as the signaling hub (primary cilia) and force generator to create extracellular fluid flow (motile cilia) during development. Dysfunction of cilia can manifest into a constellation of defects like polycystic kidney disease^1,2^, congenital heart disease^1^, primary ciliary dyskinesia^2^, retinal degeneration, etc., known as ciliopathies^3^. Nevertheless, how cells assemble a cilium remains a central outstanding question in the field.

Cilium is an extracellular hairlike organelle composed of a microtubule-based cytoskeleton called the ciliary axoneme covered with a membrane^4^. Cilium is anchored to the cell via a basal body (modified centriole) and enclosed by a specialized ciliary membrane^4,5^. The ciliary axoneme consists of the ciliary shaft and ciliary tip. The ciliary shaft is the main body of the axoneme and is made of 9+0 or 9+2 microtubule doublets in sensory or motile cilia, respectively^4^. The structure of the ciliary tip can vary across species and ciliary types; however, it is usually made of microtubule singlets in motile cilia^6,7^. The transition zone (TZ) is a structural junction between the basal body and the axoneme proposed to form a molecular sieve-like barrier^8,9^. It acts like a “ciliary gate” regulating intraciliary protein traffic^8,10,11^. To understand how cells construct such a complex structure, it is necessary to analyze the spatial and temporal order of the assembly of each component in the context of the entire cilium.

The presumption in the field is that cilia assemble bottom-up from the basal body, followed by the TZ and then the ciliary axoneme^12–14^. Ciliogenesis is initiated once the cells exit the cycling phase. The centriole now undergoes maturation to form the basal body to template ciliary growth by recruiting ciliary vesicles to the distal end. The modified centriole at this stage may build a TZ and dock to the cell membrane or dock to the cell membrane, followed by building a TZ ^15–17^. Post TZ formation, the axoneme grows by recruitment of proteins at the base of cilia by Intra-flagellar transport proteins (IFT) that ferry ciliary cargo to the tip of the cilia^18,19^.

However, in motile cilia and MCCs, more evidence is needed because cilium assembly happens in the context of other intracellular events like basal body migration and cytoskeleton remodeling^20–22^, making it challenging to tease out the spatial and temporal sequence of events. Alternatively, one can study ciliogenesis by removing mature cilia and observing the cells reassemble the entire structure without the confounding signals from basal body maturation or cytoskeleton remodeling^20,21,23^. While it is important to remember that regeneration of cilia may not be identical to *de novo* assembly, cilia regeneration studies in *Chlamydomonas reinhardtii* have provided significant insights into ciliogenesis, e.g., cargo transport, the presence of precursor pool, regulation of ciliary gene expression.^18,23,24^

The deciliation-regeneration approach offers clear advantages to understanding ciliogenesis. However, in vertebrates, this has been hindered by a few challenges. First, ciliated cells, including cells with primary cilia or multiple cilia (MCCs), are often present in tissues that are not accessible for inducing controlled deciliation and offer technical challenges to studying regeneration by imaging. The use of cell culture could circumvent this challenge. However, a detailed study in which primary cilia or MCCs have been deciliated and regenerated is absent in the literature. These obstacles can be overcome by studying MCCs of the *Xenopus* embryonic epidermis, which are external and amenable to controlled deciliation, and studying cilia regeneration by imaging ^25,26^. Previous studies have demonstrated that *Xenopus* multiciliated epithelium can be deciliated using the Ca^2+^ shock in an *in vivo* setting, and these cilia regenerate within hours ^25,27,28^. Using this *in vivo* vertebrate system of deciliation and regeneration of cilia, our study is the first to explore the fundamental questions of spatial and temporal dynamics of motile cilia assembly.

## RESULTS AND DISCUSSION

### Cilia regeneration on *Xenopus* mucociliary epithelium

The MCCs of the *Xenopus* embryonic epidermis can be consistently deciliated within 10-15 seconds of exposure to a deciliation buffer, as confirmed by the loss of acetylated (Ac.) α-tubulin signal (Fig. 1A-B). The recovery of Ac. α-tubulin signal was observed within an hour, indicating the initiation of cilia regeneration, with cilia reaching wild-type length by 6 hours (Fig. 1C). Compared to biciliate *Chlamydomonas*, where both cilia can attain full length in 90 minutes, the regeneration of cilia in MCCs of *Xenopus* takes longer, raising questions about the underlying mechanisms. One possibility is that the delay in cilia regeneration in MCCs may be due to the requirement to reassemble hundreds of cilia instead of just two in *Chlamydomonas*. Alternatively, the entire MCC may be replaced by a stem cell population rather than regenerating cilia.

**Figure 1:**
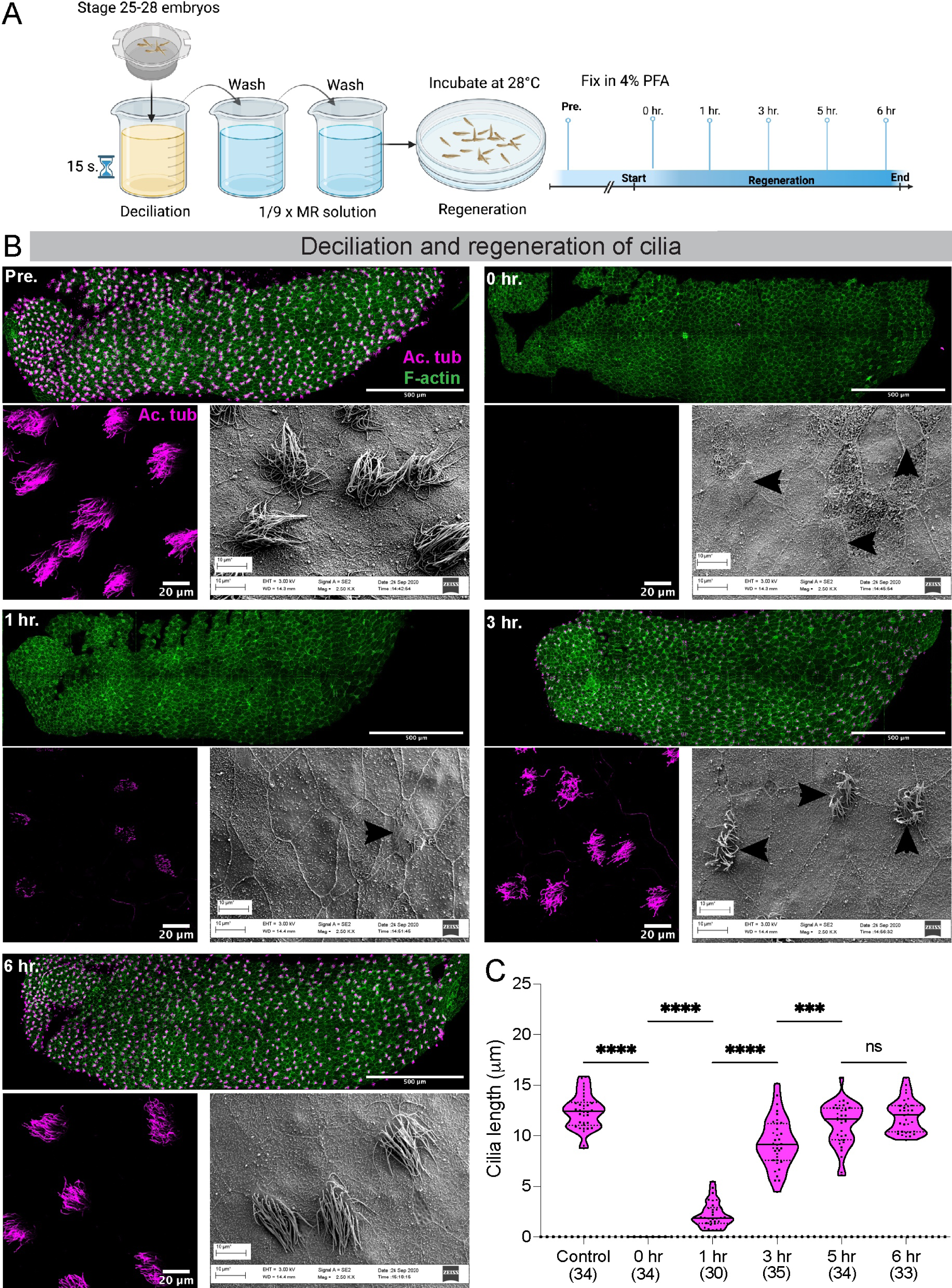
Deciliation and regeneration of cilia from the mucociliary epithelium of *Xenopus tropicalis*. A) Cartoon depiction of the deciliation and regeneration protocol of *X. tropicalis* embryos. B) Pre-deciliated embryos (Pre.) and embryos during regeneration (0 hr.,1 hr., 3 hrs., 6 hrs.) stained for cilia (Ac. α-tubulin in magenta) and F-actin (phalloidin in green). For each time point, the top panel shows the entire embryo, the lower left panel is a zoomed image of the multiciliated epithelium, and the lower right panel is an SEM image of the embryonic epithelium. Black arrows in the SEM image point to MCCs. C) Cilia length pre- and during regeneration. The values in parenthesis indicate the number of cilia measured from three trials using 10 embryos. **** - p<0.0001, *** - p<0.001, ns - not significant, ANOVA analysis was followed by Tukey’s multiple comparison test.

The mucociliary epithelium of the *Xenopus* embryonic epidermis is similar in structure and function to the mammalian airway. Damage to cilia or MCCs in the mammalian airway is suggested to be repaired by replacing the damaged MCCs using basal stem cells^29,30^. Regeneration of cilia *per se* in mammalian airway MCCs has not yet been reported in the literature. Therefore, we investigated whether *Xenopus* MCCs regenerate cilia or depend on stem cell-based replacement of damaged MCCs.

We performed live imaging of cilia regeneration to distinguish between the two hypotheses. We used *Xenopus* stem cell explants (animal caps), which auto-differentiate into an embryonic multiciliated epidermis. We harvested animal caps from *Xenopus* embryos injected with membrane RFP (to label cell and ciliary membrane) and grew them on fibronectin-coated slides. We deciliated the animal caps using the same protocol as the embryos and immediately began imaging MCCs using confocal microscopy. All MCCs showed a complete absence of cilia, indicating successful deciliation. Approximately 15 minutes post-deciliation, we could observe membrane RFP signal corresponding to regenerating cilia in MCCs. MCC specification, differentiation, radial intercalation, basal body amplification, migration, docking, and cilia formation from basal stem cells in *Xenopus* epithelium can take several hours (Personal observation). Thus, our results indicate that *Xenopus* multiciliated epithelium regenerates cilia in the same MCCs and does not undergo stem cell-based renewal of damaged MCCs. (Fig. S1A; Supplement Video 1).

### Deciliation affects the F-actin network but does not affect basal body number and polarity

Next, we investigated whether the deciliation affects only the ciliary axoneme or cilia-associated structures, such as the apical F-actin network, basal body number, and polarity. Our results show that deciliation affected the apical F-actin network significantly (calculated as cortical and medial actin) (Fig. S1B). We used Chibby or Centrin to label basal bodies and Clamp to label rootlets to assess the effect of deciliation on basal body number and polarity ^26^. Our results show that deciliation does not affect basal body number and polarity. (Fig. S1C, D).

### The Transition zone (TZ) is removed by deciliation

Next, we determined the location where the deciliation treatment severed cilia. Unicellular models such as *Chlamydomonas* and *Tetrahymena* lose cilia distal to the TZ and below the central pair (CP) microtubules^10,31–33^. Therefore, we hypothesized that *Xenopus* MCCs adopt a similar mechanism and would lose the ciliary axonemes distal to the TZ. To test the hypothesis, we labeled the TZ with an antibody to B9d1, a bona fide TZ protein^11,34–37^. To our surprise, B9d1 was lost from the deciliated MCCs (Fig. 2A, 0 hr. timepoint), suggesting that TZ was removed along with the ciliary axonemes. We did Transmission Electron Microscopy (TEM) to visualize the TZ and validate this unexpected result. TZ can be seen as an electron-dense H-shaped structure above the basal bodies ^10,38–42^. In the deciliated embryos, we could see the basal bodies; however, the H-shaped structure corresponding to the TZ was lost entirely (Fig. 2B, 0 mins. timepoint). Taken together, these data demonstrate that, unlike unicellular organisms, deciliation in *Xenopus* MCCs removes the TZ with the ciliary axoneme.

**Figure 2:**
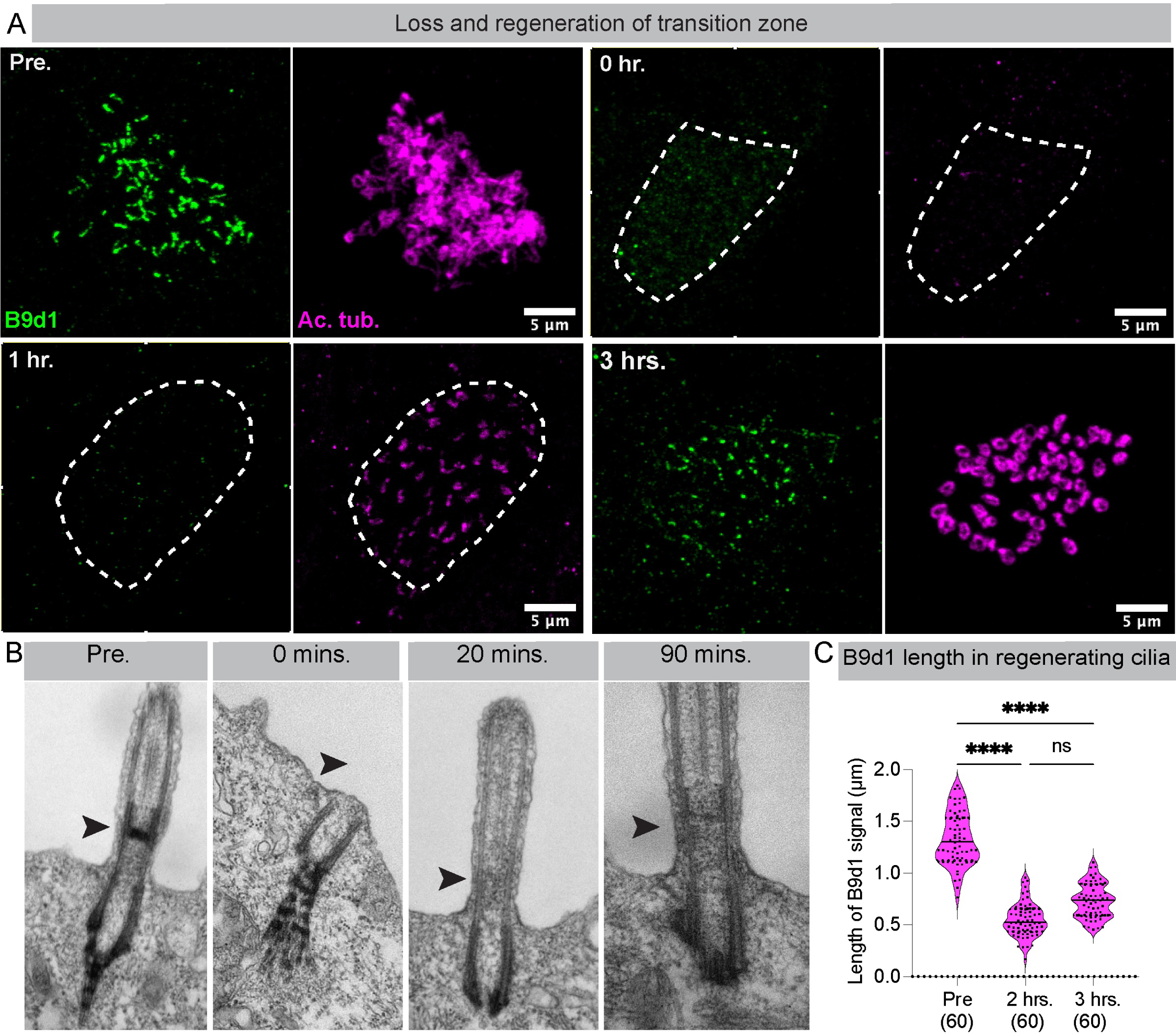
Loss and regeneration of transition zone. A) MCCs labeled with B9d1 antibody (TZ, green) and Ac. α-tubulin (cilia, magenta) in embryos pre- and during cilia regeneration (0 hr.,1 hr., 3 hrs.). MCCs are depicted by a white dashed outline. B) Representative TEM images of cilia pre-, and during the regeneration (0 mins, 20 mins, and 90 mins). The TZ can be seen as an ‘H’ shaped structure between the axoneme and basal body (indicated by a black arrow). Post-deciliation samples show a complete loss of TZ. TZ is absent after 20 mins of cilia regeneration. Premature electron-dense TZ structure can be seen after 90 minutes of cilia regeneration. C) The length of the B9d1 signal was measured pre- and during cilia regeneration (at 2 hrs. and 3 hrs.) The B9d1 signal was absent at earlier time points (0 hr. and 1 hr.). The values in parenthesis indicate the number of cilia measured from three trials using 10 embryos. **** - p<0.0001; ns - not significant, One-way ANOVA analysis followed by Tukey’s multiple comparison test.

While the TZ is lost with deciliation, the actual excision may happen at two distinct locations, distal to the basal body (between the basal body and TZ) and distal to the TZ (between TZ and CP), separately releasing the TZ and the axoneme into the medium^31,32,42^. Alternatively, deciliation may happen only distal to the basal bodies, in which case the TZ stays with the ciliary axoneme^10,31,40,42^. A previous proteomic study where authors deciliated *Xenopus* embryonic multiciliated epithelium found TZ proteins in their ciliary axoneme preparation, suggesting that the TZ is removed with the ciliary axonemes in *Xenopus* MCCs, supporting the later hypothesis ^28^. We examined whether the B9d1 signal was associated with cilia post-deciliation to test the hypothesis. To that end, we performed an *in-situ* deciliation of embryos using deciliation buffer on poly-L-lysine coated coverslips. Once removed, cilia stick to the coverslip, stained for B9d1 and Ac. α-tubulin (Fig. S2A, B). Surprisingly, we did not see any B9d1 signal with cilia (Fig. S2B, chemical deciliation). Notably, B9d1 is a soluble protein closely associated with the membrane-associated MKS protein complex in the TZ ^9,34,43^. We examined ciliary axonemes with EM to examine if the detergent in the deciliation buffer is stripping the ciliary membrane and leading to the loss of the B9d1 signal. As expected, we could not observe any ciliary membrane (Fig. S2D) supporting our hypothesis that loss of B9d1 is likely the result of loss of ciliary membrane. Therefore, to understand if the TZ remains associated with the axoneme after deciliation, we adopted an alternative approach of mechanical deciliation^44^ (Fig. S2A). The advantage of this approach is that we did not use any detergent. We observed that cilia removed from the embryo consistently showed a B9d1 signal (Fig. S2C). Thus, we demonstrate that the site of deciliation in *Xenopus* cilia is distal to the basal body, and the TZ remains attached to the ciliary axoneme.

### TZ is dispensable for the initiation of cilia assembly during regeneration

Removal of the TZ during deciliation gave us a unique opportunity not afforded by any other vertebrate or invertebrate model to study the temporal relationship between TZ and axoneme assembly. The TZ is thought to form a molecular sieve-like barrier and serve as a “ciliary gate,” regulating protein traffic into and out of the cilium^8,9^. Mutations in or knock-down/out of TZ proteins impair cilia assembly and function^11,34,36,40,45,46^. Further, recent studies have shown that Intraflagellar transport (IFT) trains assemble at the TZ, suggesting that TZ is essential for ciliary assembly and is predicted to precede ciliary axoneme assembly^10,22^. However, no direct evidence to support this hypothesis has been published.

Using our unique cilia regeneration model, we directly tested this hypothesis. We examined the restoration of the B9d1 signal as a marker to track the reassembly of TZ during the first three hours of cilia regeneration. As expected, both the B9d1 and Ac. α-tubulin signals were completely lost after deciliation. One hour into regeneration, we were surprised to see the recovery of the Ac. α-tubulin but no detectable B9d1 signal. The B9d1 signal began to appear after 2 hours of regeneration and further increased at 3 hours. (Fig. 2A). We quantified the length of the recovered B9d1 signal in the MCCs. In control embryos (pre-deciliation), the length of the B9d1 signal was 1.33 μM (± 0.33, SEM); this signal increased from no signal at 1 hour to 0.55 μM (± 0.16, SEM) at 2 hours and 0.73 μM (±0.021, SEM) at 3 hours after deciliation (Fig. 2C). This result was unexpected because, contrary to the previous speculation, cells appeared to initiate ciliary axoneme assembly in the absence of B9D1.

One limitation of our results is that we have used only one protein, B9d1, to track the loss and reassembly of TZ. We attempted to localize multiple TZ proteins in *Xenopus,* but our attempts were unsuccessful as antibodies available to *Xenopus* are particularly limited. To overcome this shortcoming, we performed electron microscopy to directly analyze the early axoneme and TZ assembly steps during regeneration. Specifically, we performed the TEM on samples taken at 0, 20, and 90 minutes after deciliation (Fig. 2B). At 20 minutes after deciliation, a small ciliary axoneme was present; however, the H-shaped structure indicating the presence of mature TZ was not yet visible. The H-shaped structure of the TZ began to appear 90 minutes post-deciliation. The TEM data supported the results from our B9d1 immunofluorescence experiments. However, one of the limitations of the TEM approach is that the sections are very thin (∼75nm), and we could miss the H-shaped structure when looking at a single section. To address this concern, we performed electron tomography of regenerating cilia at different time points (Pre., 0, 20 mins, 1 hr., 3 hrs., and 6 hrs.) to examine the timing of TZ assembly (Supplementary Videos 2-5). Consistent with the TEM data, electron tomography results demonstrated that MCCs could initiate ciliary axoneme assembly without mature TZ. The TZ was absent from the 20-minute tomogram and assembled between 1 and 3 hours after deciliation; by 6 hours, the TZ looked indistinguishable from controls. Our results suggest that the cells can incorporate TZ (or components of TZ) once the axoneme is assembled, suggesting a dynamic nature of TZ proteins. An earlier study in *Chlamydomonas* also reports the dynamic nature of a TZ protein, CEP290, incorporated into the pre-assembled axoneme^31^. During the mating process, there was an exchange of CEP290 from the wild-type cilia to the cilia of the CEP290 deficient mutant during the mating process ^31^. While the previous study showed one TZ protein, CEP290, to be dynamically assembled in the cilia, and our study shows the incorporation of another TZ protein, B9D1, we speculate that this could apply to many TZ proteins where they could be dynamically assembled in cilia.

The structure of the ciliary axoneme is different at the proximal end, where the TZ is located, and the distal end, where the ciliary tip is located. Like the TZ, the ciliary tip comprises of a distinct protein complex of unknown function^6,7,39,47,48^. Since the TZ showed a delayed regeneration, we asked if the ciliary tip proteins assembled after the ciliary shaft was built. To test this speculation, we used mScarlet-Sentan^49^ and RFP-Clamp ^50^ proteins known to localize to the tip of motile cilia. As reported earlier^49^, we observed that mScarlet-Sentan and RFP-Clamp localized to ciliary tips in control *Xenopus* MCCs (Fig. 3A, S3A). After deciliation, ciliary Sentan and Clamp signals were lost as expected. However, when regeneration began, we noticed that small stubs of ciliary axonemes at 1 hr. post-deciliation were decorated with the Sentan and Clamp signals (Fig. 3A, B and S3A, B). We measured both the Clamp and Sentan signals at different time points during regeneration. We observed that, like pre-deciliation controls, Sentan and Clamp signals became stronger at the ciliary tips as regeneration progressed. (Fig. 3B, S3B). Our data suggests that ciliary tip proteins like Sentan and Clamp are trafficked immediately into regenerating cilia and gradually begin to concentrate at the tips.

**Figure 3:**
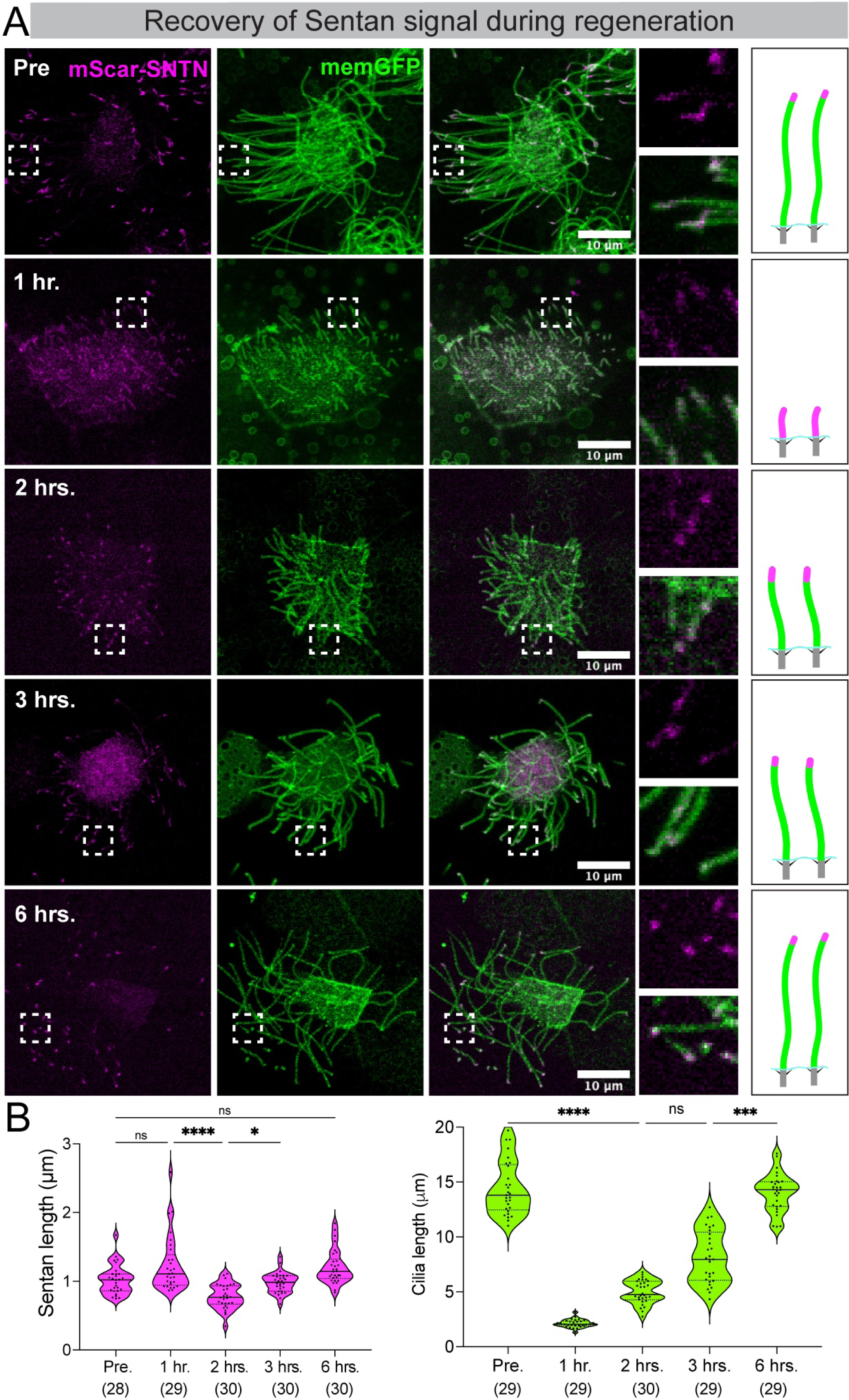
Ciliary tip protein Sentan is localized to the ciliary axoneme during the early stages of regeneration. A) MCCs are labeled with mScarlet-Sentan decorating ciliary tips (magenta) and membrane-GFP (cilia, green) at various stages of cilia regeneration. Sentan can be seen at the tips of cilia in pre-deciliated samples. After 1 hr. post-deciliation, Sentan spans the entire length of ciliary axonemes. At 2 hrs., Sentan starts accumulating at the ciliary tips. At 3 and 6 hr. Sentan signal appears similar to pre-deciliated samples. B) The Sentan signal length (left panel) and cilia length (right panel) were measured and compared among different time points. The values in parenthesis indicate the number of cilia measured from three trials using 9 embryos. **** - p<0.0001; *** - p<0.0005; * - p<0.05 ns - not significant, Kruskal-Wallis test, followed by Dunn’s test.

### TZ protein B9d1 is newly synthesized during cilia regeneration

We next addressed why there was a delay in assembling TZ. The clue came from an earlier study where photobleaching of B9D1 and some other TZ proteins (in a pre-existing TZ) did not recover to pre-bleaching levels^51^. This study indicated that unlike some proteins needed for axoneme maintenance, e.g., IFTs^18^, the TZ proteins such as B9d1 may not be needed for cilia maintenance and thus are not present in the cytosolic pool. Consistently, we did not observe any B9d1 signal near the base of the cilia or in the cytoplasm. Therefore, we hypothesized that post-deciliation MCCs newly synthesize TZ proteins, such as B9D1, resulting in delayed assembly of TZ.

To test this hypothesis, we treated deciliated embryos with either vehicle (DMSO) alone or Cycloheximide (CHX), a protein synthesis inhibitor. We collected embryos every hour for the first 3 hours and quantified the B9d1 signal in the regenerating cilia (Fig. 4). By 3 hours, control MCCs showed a significant increase in the B9d1 signal. However, treatment with CHX completely abolished B9d1 recovery. These results suggest that B9d1 is absent in the ciliary precursor pool and requires new transcription/translation during regeneration.

**Figure 4:**
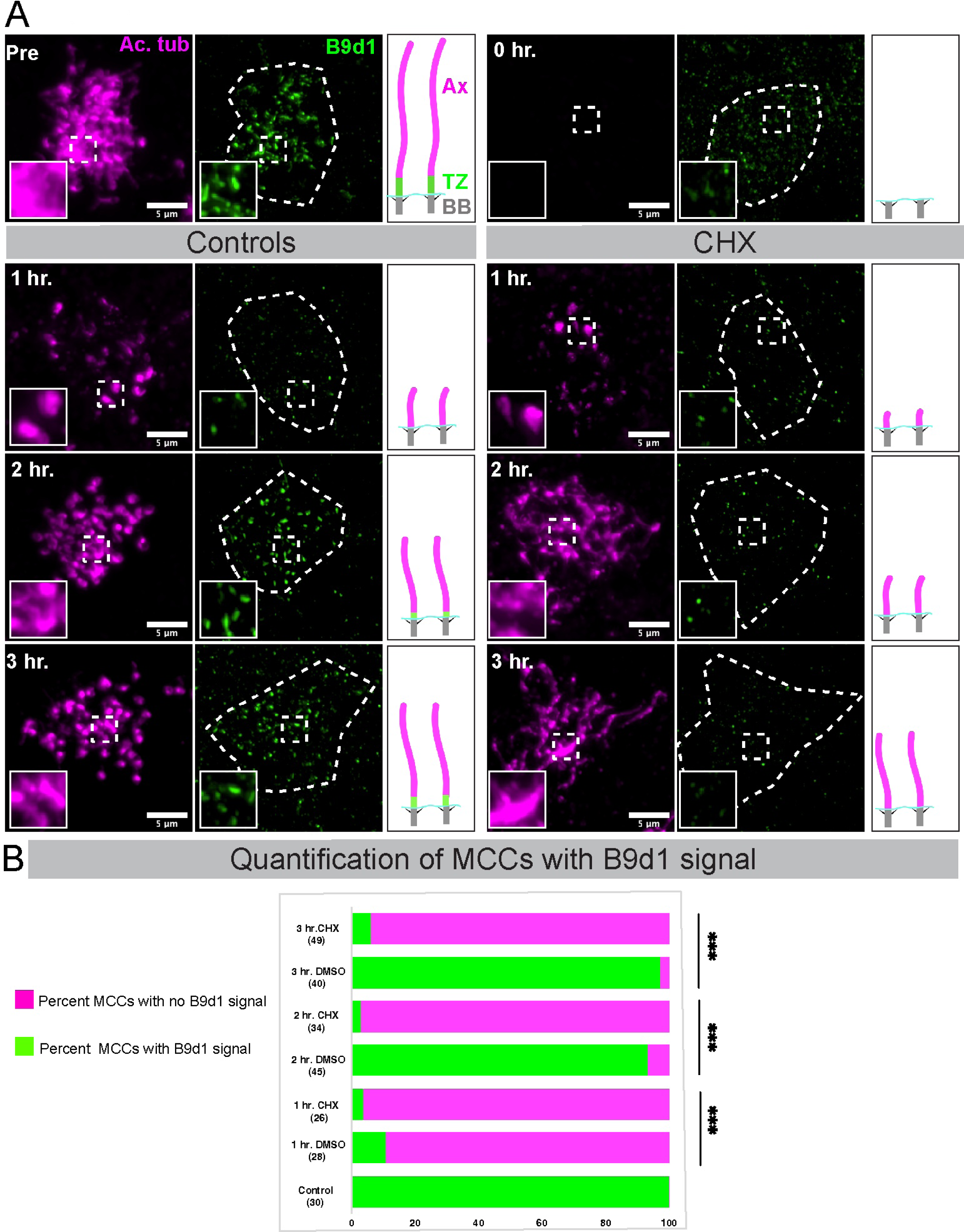
TZ protein B9d1 requires new protein synthesis during regeneration. A) Embryos treated with DMSO alone (Vehicle) or cycloheximide (CHX) in DMSO were stained for B9d1 (green, TZ) and Ac. α-Tubulin (magenta, cilia) that were collected pre-deciliation and during cilia regeneration (0 hr., 1 hr., 2 hrs., 3 hrs.). The cell outline is marked with a white dashed line, and the dashed square box in the B9d1 channel is zoomed and depicted in the inset. While cilia can regenerate in both treatments (controls and CHX), the B9D1 signal is recovered only in controls and remains completely absent in the CHX-treated samples at 2 hrs. And 3 hrs. timepoints. Ax: Axoneme, TZ: transition zone, BB: Basal body B) The percent of MCCs positive for the B9d1 signal (green) and the absent B9d1 signal (magenta). Numbers in parentheses indicate the number of MCCs counted from three trials using 9 embryos. *** - p<0.001, chi-square test using VassarStats to test the significance.

### Synthesis of a new pool of ciliary proteins is required to support cilia regeneration

Our results from the previous experiments raised the question: does deciliation trigger new protein synthesis in MCCs? From the live imaging experiments, we noted that cilia regeneration begins within approximately 12-15 minutes after deciliation. *Chlamydomonas* also begin cilia reassembly within minutes after deciliation. Studies have shown that *Chlamydomonas* uses the precursor pool of proteins for rapid cilia reassembly; however, this precursor pool is insufficient to assemble full-length cilia ^52,53^. Loss of cilia triggers the synthesis of new mRNA and proteins to support subsequent cilia assembly ^53^. In contrast to *Chlamydomonas*, sea urchins contain excess proteins that can support full regeneration of cilia up to four times ^54^. Therefore, we speculated that *Xenopus* MCCs might have a precursor pool of unassembled proteins that allow the cell to begin cilia reassembly quickly. However, it remains unknown whether *Xenopus* MCCs require new transcription and translation of some/all proteins to complete cilia regeneration or whether the unassembled ciliary protein pool is enough to build all cilia.

To test this hypothesis, we blocked protein synthesis immediately post-deciliation using Cycloheximide (CHX); thus, the precursor pool is the only source for building new cilia. (Fig. 5A). *Chlamydomonas* has only two cilia, but in *X. tropicalis* MCCs, there are ∼150 cilia. Therefore, CHX treatment could result in either the cell having a sufficient protein pool to regenerate all cilia comparable to vehicle control or a limited pool that results in MCCs choosing between the number and length of cilia.

**Figure 5:**
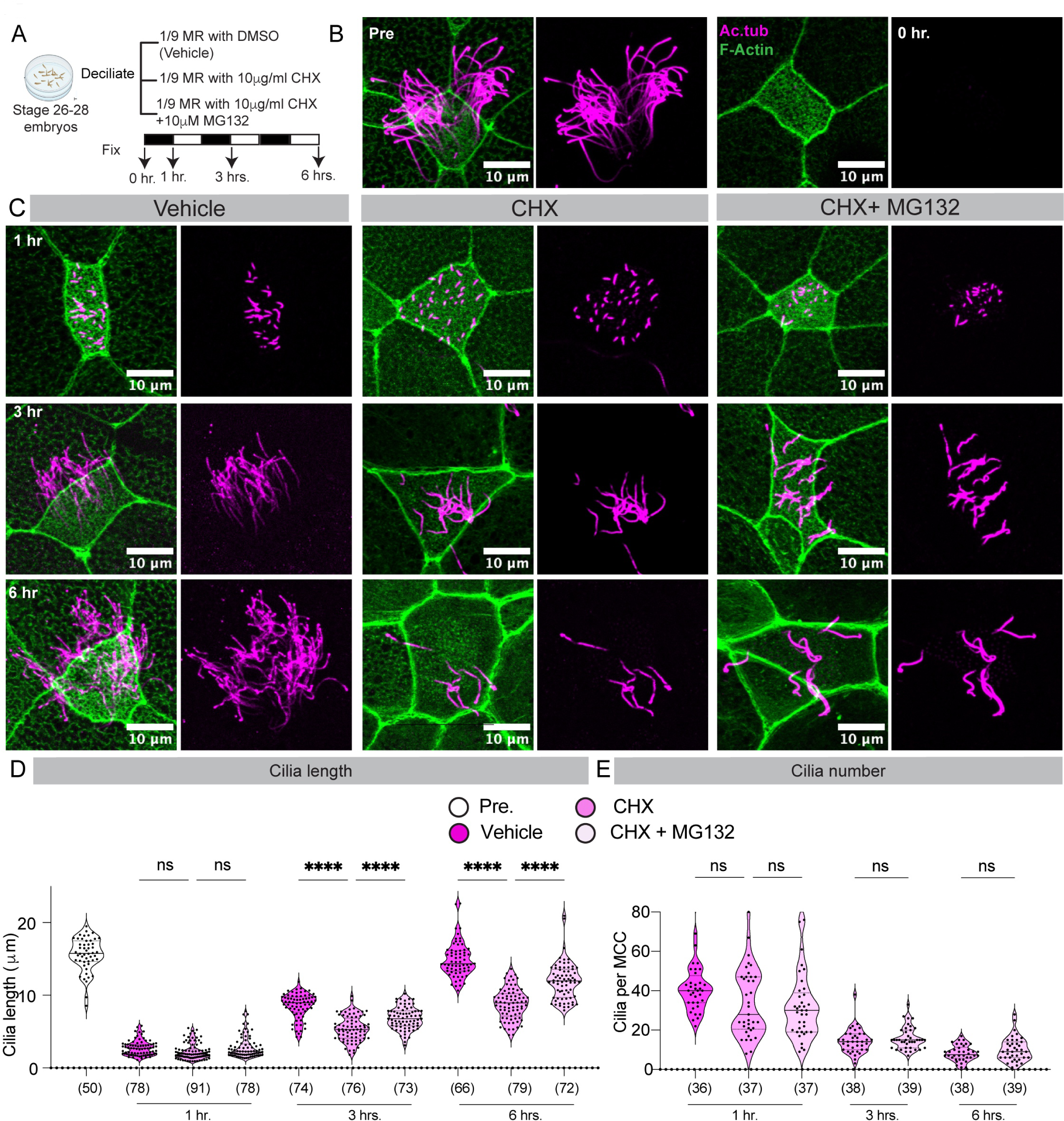
Complete cilia regeneration requires new protein synthesis. A) Deciliated embryos were treated with DMSO (Vehicle), cycloheximide (CHX), and CHX with MG132 in DMSO and collected during different time points of cilia regeneration (0 hr., 1hr., 3 hrs., 6 hrs.). B) Pre-deciliated and 0 hr. deciliated MCCs stained for with Ac. α-tubulin antibody (cilia, magenta) and Phalloidin (F-actin, green). C) MCCs treated with vehicle, CHX, and CHX+MG132 at 1hr., 3hrs., and 6 hrs. D) Graph showing the cilia lengths at different times pre-deciliation and during cilia regeneration in Vehicle, CHX, and CHX+MG132. The number in parenthesis indicates the number of cilia counted from three trials using 9-10 embryos. ****- p<0.0001, ns = not significant, Kruskal Wallis test, followed by Dunn’s multiple comparison test. E) Graph showing the effect of CHX and CHX with MG132 treatments on cilia number per MCC during regeneration. The number in parenthesis indicates the number of MCCs counted from three trials using 9-10 embryos. ****- p<0.0001, **- p<0.01, ns = not significant, Kruskal Wallis test, followed by Dunn’s multiple comparison test.

We observed that embryos treated with CHX showed shorter cilia than controls during regeneration (Fig. 5B, C, D, S4A, B). Specifically, there was no significant difference in the ciliary length in the first hour between control and CHX treatment (vehicle vs. CHX, 2.75±1.04μm vs. 2.24±1.12μm, mean±SD, Fig. 5D, and S4A and B). However, cilia did not grow to the same length in CHX treatment as vehicle treatment at 3 and 6 hrs. (3 hrs. vehicle vs. CHX, 8.53±1.74 μm, vs. 5.40±1.73 μm, and 6 hrs. vehicle vs. CHX, 14.87±2.38μm vs. 8.8± 2.14μm; mean±SD; Fig. 5D and Fig. S4A and B). Thus, ciliary assembly is compromised during regeneration with CHX treatment.

Next, we measured the number of cilia per MCC during cilia regeneration. While we could count the cilia number at 1 hr. in both treatments (1 hr. vehicle 41.61±12.71 vs CHX; 33.78±17.27 cilia/MCC, Fig. 5C, S4C), it was challenging to count the cilia in vehicle-treated samples after 3hrs. and 6hrs. due to the long cilia that clumped/entangled during the staining process. However, the basal body number did not change with the vehicle and CHX treatment (Fig S5A and B). Assuming all the basal bodies can assemble cilia, we can estimate 125-150 cilia/MCC in vehicle treatment (Fig S5B). In contrast to the controls, we noted that the cilia number at 1 hr. time point in CHX-treated MCCs was higher than the 3 hrs. (1 hr. CHX vs. 3 hrs. CHX; 33.78±17.27 vs. 14±6 cilia/ MCC, Fig. 5B, E, S4C). This number was further reduced by the end of 6 hrs. (8 ± 4 cilia/MCC; mean±SD, Fig. 5C, E, Fig S4C), suggesting that the CHX treatment reduced the cilia number without affecting the basal body number.

We speculated that this dramatic reduction in cilia number and length in CHX treatment could be due to the blocking of new protein synthesis and the ubiquitin-dependent proteasomal degradation of the existing ciliary protein pool in the cell. To test the contribution of protein degradation on the regeneration of cilia length and number, we blocked protein degradation using the drug MG132 (Fig 5A)^55^. Treating the cells with MG132 with CHX did not significantly increase the number of cilia at any time point (Fig. 5C, E, and Fig. S4C). However, CHX+MG132 led to a significant increase in cilia length compared to CHX treatment, by an average of 3μm by 6 hours. (3 hrs. CHX vs CHX+ MG132; 5.4 ± 1.73μm vs. 6.85±1.53 μm, 6 hrs. CHX vs CHX+ MG132; 8.8 ±2.14μm vs. 11.93±2.53 μm; mean ± SD; Fig. 5C, D, S4A, B).

Overall, these experiments reveal an interesting observation: with a limited protein pool, MCCs prefer longer cilia over more cilia. We speculated that MCCs optimize the length and number of cilia to generate optimal beating force. Therefore, the reason for MCCs to prefer length over number was unclear. A longer cilium can exert more beating force over a shorter cilium up to a certain length. However, increasing length beyond a threshold can increase the hydrodynamic drag ^56,57^. However, in MCCs, this calculation is complex because both cilia length and number play important roles in generating the optimal force necessary for mechanical coupling with surrounding MCCs ^58^. To understand this relationship, we used mathematical modeling to generate a force matrix for different numbers and lengths of cilia per cell (Supplement Table 1). Our modeling data shows that the force exerted by the beating cilia increases more for a unit increase in length than the number. For example, one 12-μm-long cilium will exert higher force than 12 cilia of 1-μm length (Supplementary Table 1). When applied to the number and lengths in our data, we show that the theoretical force generated by vehicle-treated MCCs increased during regeneration. Further, as expected, it was higher than the CHX and CHX+MG132 treated MCCs (Fig. 6A, B, Supplementary Table 1). Within the CHX-treated samples (CHX and CHX+MG132), there was an increase in force generation from 1hr. to 3hrs., and 3hrs. to 6 hrs. despite a gradual decrease in the number of cilia. (Fig. S4C, Fig. 6A, B). Interestingly, the reduction in cilia number is correlated with an increase in cilia length, supporting the theoretical calculations that force generation has a quadratic relationship with cilia length compared to a linear relationship with cilia number. However, one could argue that MCCs have more cilia building blocks as a precursor pool. As a result, over 6 hrs. of regeneration, there was an increase in total cilia quantity, which led to increased force generation (cilia quantity = avg. number of cilia X avg. cilia length). Therefore, we measured total cilia quantity over time in CHX and CHX+MG132 treatments. Our data shows that total cilia quantity in CHX treatment remained unchanged (75.66μm at 1 hr., 77.16μm at 3 hrs., and 72.78μm at 6 hrs.), suggesting that the ciliary proteins were likely redistributed from shortened cilia to support fewer but longer cilia. This phenomenon was also observed in *Chlamydomonas* when one of the two flagella is removed in the presence of CHX, and the intact flagellum is shortened to support the growth of the amputated flagellum ^18,59^. On the other hand, there was a slight increase in total cilia quantity over time in CHX+MG132 treatment (89.56μm at 1 hr., 108.41μm at 3 hrs., and 122.64μm at 6 hrs.). However, this increase was minimal (21.04% from 1 hr. to 3hrs. and 13.12% from 3 hrs. to 6 hrs.) compared to the calculated force generation (214.7% from 1 hr. to 3 hrs. and 163.06% from 3 hrs. to 6 hrs.), which was more proportionate to the change in cilia length (241.7% from 1 hr. to 3 hrs. and 174.41% from 3 hrs. to 6 hrs.). Therefore, we propose that cilia length is a major determinant of force generation. While our model is based on the previously published literature ^56,57,58^, it is still a simplified model and does not consider other factors, such as the coordination of beating cilia, the mucus viscosity, and the spacing/distribution of ciliary arrays on the cell, which could affect the effective force transmission.

**Figure 6:**
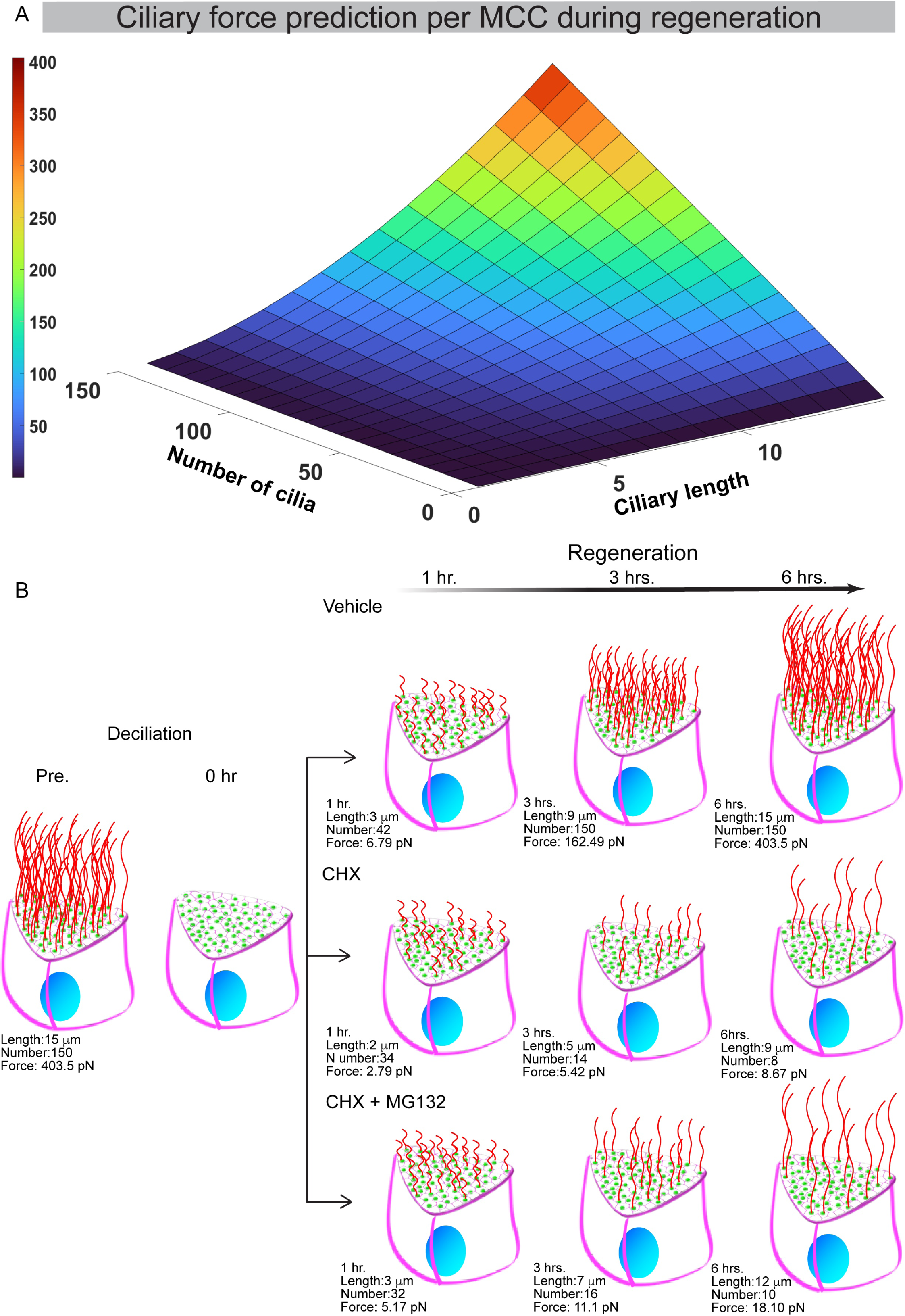
Mathematical model describing the contribution of cilia length and number to force generation in MCCs. A) The visualized 3-D plot shows a linear relationship between the number of cilia and the force generated by MCC, while a quadratic relationship exists between the length of cilia and the force generated by MCC. The individual force values for different cilia lengths and numbers are given in Supplementary Table 1. B) The cartoon depiction of the data shows that, a decrease in cilia number and an increase in cilia length were correlated with increased force generation (calculated from the model) over time in both CHX and CHX+MG132 treatments.

Our model would also benefit from a direct test to measure bead movement as a readout of force generation by these MCCs. However, there are technical challenges involved in accurately measuring the bead movement over the surface of single MCCs with different cilia lengths and numbers. Even if we overcome this challenge, we do not understand when the motility machinery is fully incorporated into a regenerating cilium. Moreover, in the CHX treatment, whether the machinery proteins are present as precursors or require new transcription/translation similar to the TZ protein B9D1. Nonetheless, mathematical modeling coupled with the experimental data on cilia length and number sheds light on an important cell biology question of how cells balance the number and size of organelles to optimize functional output.

### MCCs redistribute proteins among basal bodies to regenerate fewer but longer cilia

Given the total cilia quantity in CHX treatment remains unchanged from 1 hr. to 6 hrs., and MCCs cannot synthesize new proteins, we hypothesized that existing proteins essential for ciliogenesis accumulate at a few basal bodies to support fewer longer cilia. A similar phenomenon was observed previously in *Chlamydomonas*; when *Chlamydomonas* loses one of its two flagella, it resorbs the longer flagellum to redistribute the material to the regenerating flagella to support its assembly until both flagella reach the same length ^18,52^. However, unlike *Chlamydomonas* cells, *Xenopus* MCCs have hundreds of cilia, and basal bodies are not physically connected, raising the possibility of distinct mechanisms.

IFT proteins are necessary to assemble the axoneme by transporting the axonemal proteins to the tip of the growing structure ^19,27,60^. Ciliogenesis and cilia regeneration occur by accumulating several ciliary proteins, including IFTs at the base of cilia in *Chlamydomonas* and *Xenopus* ^10,18,27^. Therefore, we examined the pool of three IFT proteins (IFT20, IFT80, and IFT43) at the base of regenerating cilia as a proxy for essential ciliogenesis proteins required for cilia regeneration in the vehicle and CHX-treated embryos (Fig. 7, Fig. S6, S7). In pre-deciliation-and 0 hr. deciliated MCCs, all basal bodies were decorated with IFT proteins (IFT20, IFT80, and IFT43) signal, demonstrating that IFTs were similarly distributed among all basal bodies. In vehicle-treated embryos, all basal bodies remained equally decorated with IFT signal throughout regeneration as expected (Fig. 7A, S6A, S7A). In contrast, in the CHX-treated cells, we observed that IFT proteins gradually enriched at a subset of the basal bodies as regeneration progressed. Interestingly, the basal bodies that were saturated with the IFT signal were also ciliated at all time points (Fig. 7A”, B, S6A”, B, S7A”, B), suggesting that in the absence of new protein synthesis, IFT proteins enrich a few basal bodies and support the assembly of long cilia. It must be noted that the non-ciliated basal bodies at 6 hrs. possess some IFT-GFP signal (IFT20, IFT80, and IFT43); we speculate this could be due to multiple reasons. One, some minimal amount of IFT is always associated with basal bodies. Second, these IFT molecules are in the process of getting redistributed. Note that 6 hours is sufficient to regenerate full-length cilia in vehicle-treated MCCs, but we do not know if this dynamic applies to CHX treatment. Third, this results from ectopic overexpression, where an excess of proteins could be associated with non-ciliated basal bodies.

**Figure 7:**
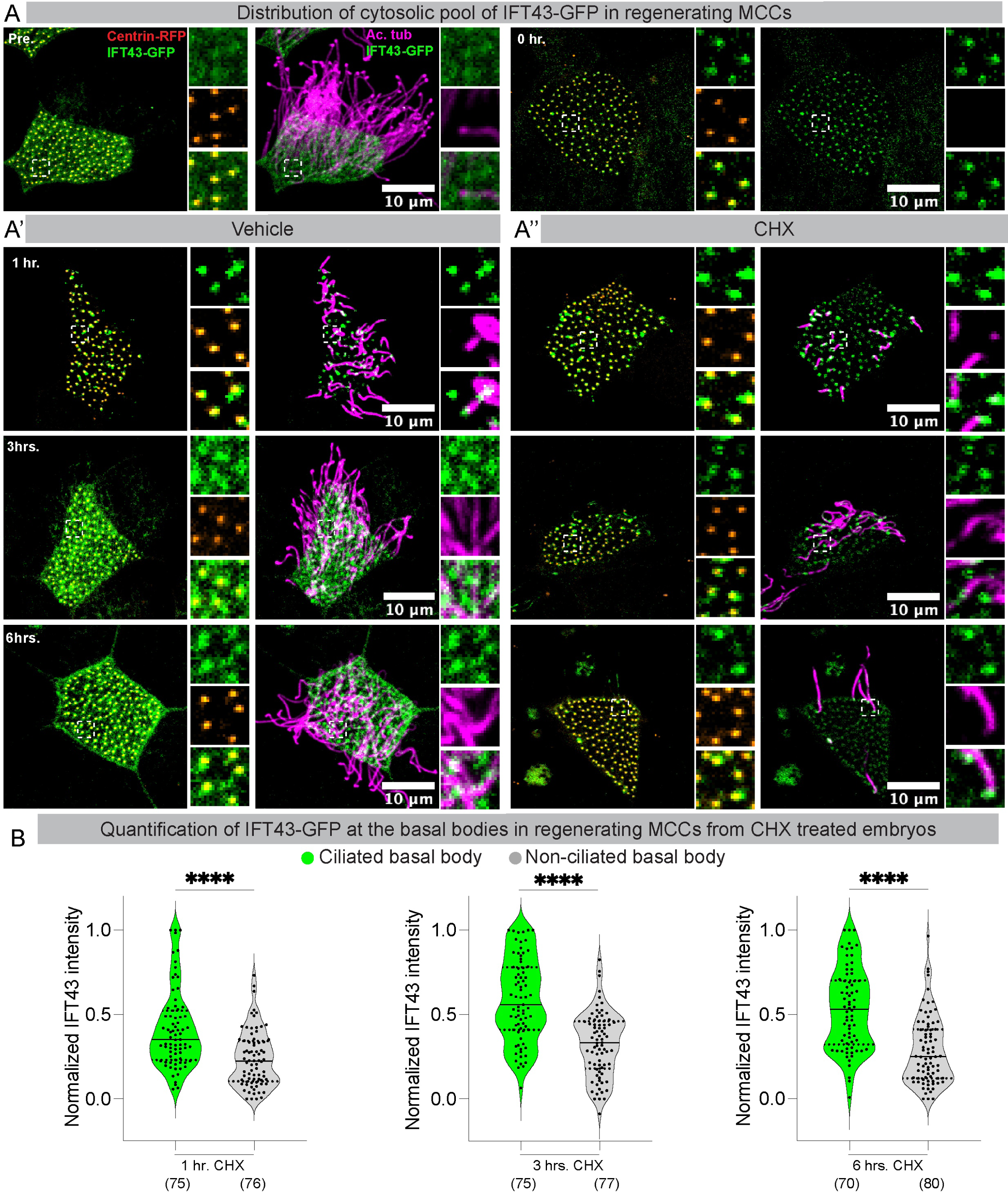
Distribution of ciliary precursor pool (IFT43-GFP) in MCCs. A) Embryos injected with IFT43-GFP (green) and Centrin-RFP (orange hot, basal bodies) were deciliated at stage 28 (0 hr.) and were split into two experiments (DMSO and CHX). A’, A’’) The embryos in both sets were allowed to regenerate cilia for 6 hours. After fixation, the embryos were stained for cilia(magenta). Note that the control MCCs at all time points (1 hr., 3 hrs., and 6 hrs.) have multiple cilia regenerating, and the IFT43-GFP intensity is uniform at every basal body. In contrast, the number of regenerating cilia decreases with time in CHX-treated samples, and the IFT43-GFP is enriched at a few ciliated basal bodies. B) The intensity of IFT43-GFP associated with ciliated (green) vs. non-ciliated (gray) basal bodies in the same MCC (in CHX-treated samples) at different time points during cilia regeneration. A total of 8-10 basal bodies per MCC (4-5 ciliated and 4-5 non-ciliated) and 5 MCCs were chosen, and the mean gray value was estimated and normalized to the maximum and the minimum values in the set. The value in parenthesis indicates the number of basal bodies analyzed (with and without IFT43-GFP) from 9 embryos from three independent trials. Note the significant difference in the IFT43-GFP signal intensity at ciliated vs. non-ciliated basal bodies at all time points. **** - p<0.0001, Kruskal Wallis test, followed by Dunn’s multiple comparison test.

While *Chlamydomonas* also redistributes the proteins between cilia to coordinate their regeneration, significant differences between the model systems raise fascinating questions about how cells sense information and make decisions that may require quantitative data ^61,62^. For example, *Chlamydomonas* has only two basal bodies that are proximally placed and physically connected. Thus, the basal bodies share the protein pool required for assembly. On the other hand, *Xenopus* MCCs nucleate hundreds of basal bodies. While they are connected via the cytoskeletal network ^20,26,63^, they are not physically connected like *Chlamydomonas*. Moreover, each cilium has its pool of proteins at the basal body to support the assembly. Given hundreds of basal bodies in *Xenopus* MCCs, how does inter-basal body transport operate? One possibility is that proteins are transported by diffusion to capture mechanism from the cytosol to cilia ^27,64^. However, another possibility is the direct transport between basal bodies using a cytoskeleton (microtubule) network^65,66^. Hibbard et al. tested this possibility by depolymerizing microtubules with nocodazole and cold shock and showed that IFT recruitment at basal bodies could be independent of cortical microtubules ^27^. However, this experiment was not carried out when the cell was experiencing a shortage of proteins, as is the case for CHX treatment. Our results also show that cells ensure that at least a few basal bodies can assemble near wild-type length cilia, raising questions about how these basal bodies are selected. One possibility is that ciliary axonemes must reach a minimum length to be selected to grow longer. This is not an issue in wild-type cells where proteins are abundant; however, in a protein-deficient state, only cilia that reached the minimum length continue assembling, whereas others resorb because of competition for the available pool of proteins.

In conclusion, our study reveals new and significant findings and raises exciting questions about the mechanisms of cilia regeneration in vertebrates. We show that the location for severing the ciliary axoneme in MCCs was distal to the basal body, which differs from unicellular organisms like *Chlamydomonas*. Most interestingly, MCCs initiate the assembly of the ciliary tip and ciliary shaft before the assembly of the TZ, after which both grow concurrently (Fig. 8). One possibility is that there is a lack of a cytosolic pool for one or more proteins that are rate-limiting for TZ assembly. These findings raise intriguing questions about the spatial-temporal sequence of cilia assembly and the importance of TZ in ciliary assembly. For example, is the TZ dispensable for motile ciliogenesis? When are other components needed for ciliary stability (e.g., microtubule inner proteins) transported into the cilium? How do MCCs decide the spatial location of the central pair microtubules without TZ? Moreover, the study also highlights the importance of new transcription/translation in completing cilia regeneration. Without protein synthesis during cilia regeneration, MCCs produce few longer-length cilia over multiple short-length cilia. Our mathematical model suggests that this optimization helps MCCs exert more force per cell. This study also provides important insights into the cellular decision-making process that regulates the size and number of organelles to maximize their function.

**Figure 8:**
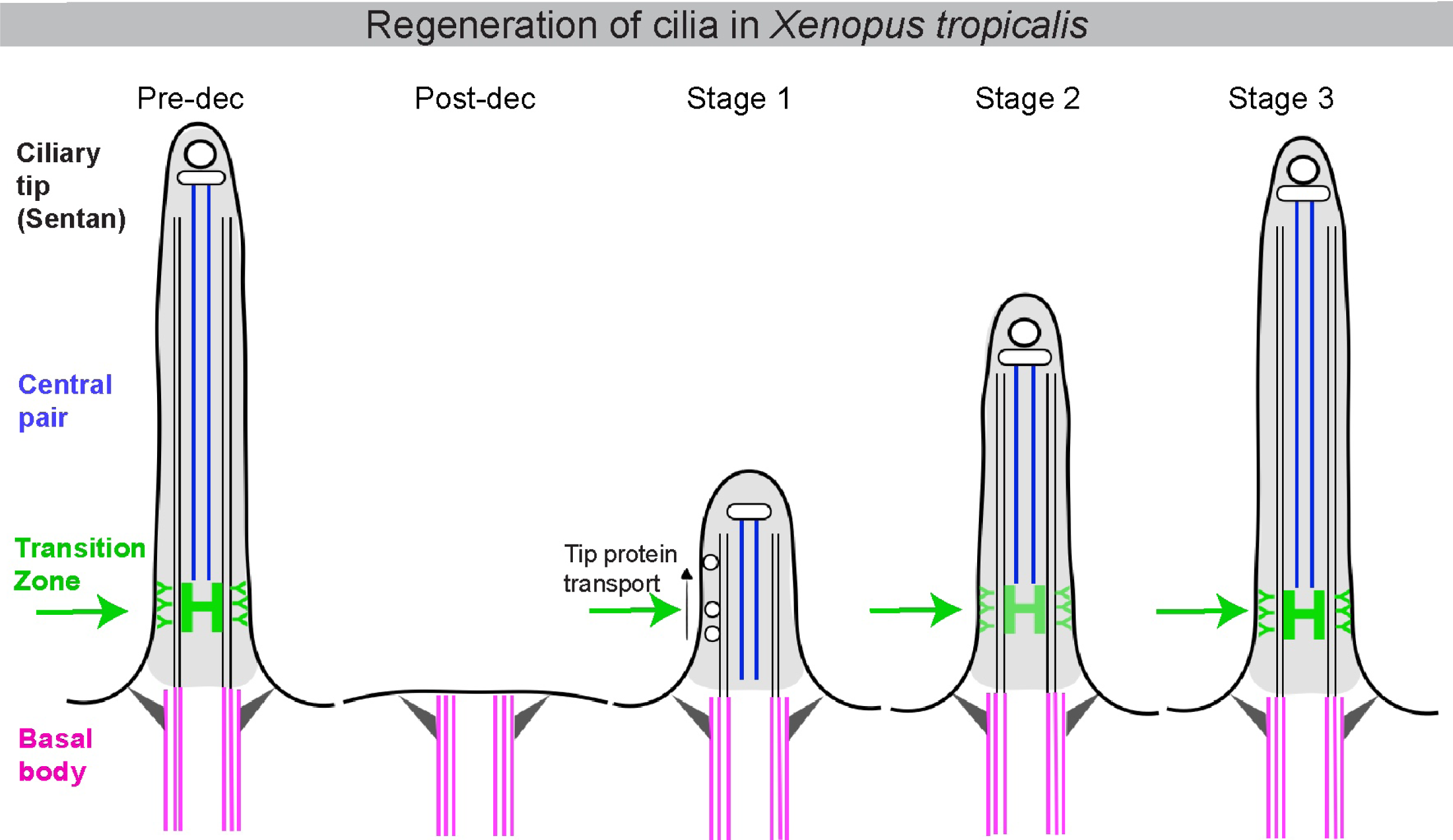
Model for cilia regeneration in *Xenopus tropicalis* embryos. Cilia regeneration in *Xenopus tropicalis* embryonic MCCs is multi-staged and is depicted in the cartoon. Post-deciliation, the TZ is lost with cilia. During stage 1 of regeneration (0-2 hours), the cilia grow without the TZ. Also, the cilia tip proteins Sentan and Clamp are immediately trafficked into the ciliary axoneme. By stage 2 (2-4 hours), a partial transition zone appears, and ciliary tip proteins begin to accumulate at the distal end of the axoneme. The cilia look like pre-deciliated samples by the end of regeneration.

## MATERIALS AND METHODS

### Animal husbandry and *in-vitro* fertilization

Frogs (*Xenopus tropicalis*) were bred and housed in a vivarium using protocols approved by the University of Virginia Institutional Animal Care and Use Committee (IACUC). Embryos needed for experiments were generated using *in-vitro* fertilization as described before ^26,67^. Briefly, the testes from male frogs were crushed in 1x MBS (pH-7.4) with 0.2% BSA and added to eggs obtained from the female frogs. After 3 minutes of incubation, freshly made 0.1x MBS (pH-7.8) was added, and the eggs were incubated for 10 more minutes till contraction of the animal pole of the eggs was visible. The jelly coat was removed using 3% cysteine in 1/9 x MR solution (pH 7.8-8.0) for 6 minutes. The embryos were either microinjected with DNA/RNA and raised at 25°C or 28°C till they reached appropriate stages for fixation and staining or mucociliary organoid (animal cap) dissection. Embryos were staged as described previously ^68^.

### Animal cap / mucociliary organoid dissection

Embryos at stage 10 were used for animal cap dissection. The animal pole from the stage 10 embryos was dissected using a fine hair tool in a pool of Danilchik’s for Amy (DFA) medium. The dissected tissue was trimmed to remove the equatorial cells from the sides. This tissue was then laid on a coverslip or glass slide coated with fibronectin (Sigma Cat No. F1141) immersed in DFA solution. The tissue was allowed to grow and spread on the fibronectin-coated slides/coverslips overnight till they reached stage 28.

### DNA, RNA, and microinjections

Plasmids used in this study are listed in the key resource table. For generating Chibby-BFP, the full-length Chibby gene was cloned to the N-terminus of mtagBFP2 in the pCS2+ vector using Gibson assembly. RNA used in this study was generated by linearizing the plasmids and *in-vitro* transcribed using the mMessage and mMachine SP6 transcription kit (Thermo Fisher Scientific). Microinjections were done using glass needles using the Pico-liter microinjection system (Warner instruments). All the microinjections were done at the 1-cell stage except for Figure 2, where the membrane RFP was injected in 1 of the four blastomeres at the four-cell stage.

### Chemical deciliation of embryos and mucociliary organoids and drug treatments

Embryos, after they reach stage 28, were used for deciliation experiments. A basket was designed for deciliation by cutting a 50 ml centrifuge tube. One of the open ends of the tube was heat-sealed with a muslin cloth. A 6-well plate was filled with deciliation solution (75 mM CaCl_2_, 0.02% NP40 in 1/3 MR solution). The rest of the five wells were filled with 1/9x MR. Anesthetized embryos were transferred to this container and dipped in a deciliation solution (75 mM CaCl_2_, 0.02% NP40 alternative in 1/3 MR solution) in a 6-well plate for 10-15 seconds. Post-deciliation, embryos with the container were dipped/washed with 1/9x MR in the rest of the wells of the 6-well plate. The embryos were then transferred to a petri dish with 1/9x MR with gentamicin and incubated at 28°C. Embryos were collected during various stages of cilia regeneration, as mentioned in the text and figure legends. For animal caps, the glass slide/coverslip was removed from the DFA medium and dipped in the deciliation solution for 15 seconds. Post-deciliation, the glass side/coverslip was dipped in 1/9x MR for 30 seconds and transferred to DFA until the cilia were regenerated to the appropriate stage. Control or deciliated embryos were transferred to 1/9 X MR solution containing 10 mg/ml of cycloheximide (CHX) or DMSO for cycloheximide experiments. MG132 was used to prevent ubiquitin-mediated protein degradation; we used 10 μM of MG132 dissolved in DMSO for treatment post-regeneration.

### Mechanical deciliation

Poly-L-lysine-coated cover glass was used for the mechanical deciliation of embryos. 25-30 embryos at NF stage 28-30 were transferred to the poly-L-lysine-coated cover glass, and any residual 1/9 x MR was removed. A mechanical pressure was applied by a glass slide, placing it on top of the embryos for a minute. After a minute, the embryos were discarded, and the cilia stuck to the coverglass were fixed in chilled methanol for 10 minutes. And rehydrated using 1x PBS, followed by blocking and staining with anti-Ac. α tubulin antibody and anti-B9D1 antibody.

### Immunofluorescence staining and imaging

The *X. tropicalis* embryos used for the study were fixed post-stage 28 with appropriate fixative agents. For most experiments, 4% paraformaldehyde (PFA) was used as a fixative, and 100% chilled methanol was used to fix the embryos for anti-B9d1 staining. After fixation, the embryos were washed three times with PBST (1x PBS with 0.2% Triton X-100) for 10 mins each and then incubated in a blocking solution (3% BSA in PBST) for 1 hour. Appropriate antibody was added to the embryos, incubated for 1 hour at room temperature, and rewashed three times for 10 mins each with PBST. A conjugated secondary antibody was used to stain embryos for 1 hour. The embryos were washed thrice with PBST, stained with phalloidin in PBST for 45 minutes, and washed once post-staining. The embryos were mounted and imaged. Confocal imaging was performed using the Leica DMi8 SP8 microscope with a 40x oil immersion objective (1.3 NA). Images were captured at 1x or 4x zoom and adjusted (brightness and contrast), analyzed, cropped in Fiji, and assembled in Adobe Illustrator software.

### Sample preparation for electron tomography

*Xenopus* stage 26-28 embryos Pre and post deciliation were fixed in 2.5% glutaraldehyde and 2% paraformaldehyde in 0.1M sodium cacodylate buffer pH-7.4 for 1 hour, rinsed in the buffer, then post-fixed in 1% osmium tetroxide. Subsequently, embryos were rinsed in buffer and stained in 2% aqueous uranyl acetate for another hour. Embryos were rinsed and dehydrated in an ethanol series (15min 30% ethanol, 15min 50% ethanol, 15min 70% ethanol, 15min 90% ethanol, 3x 30min 100% ethanol). Dehydrated embryos were infiltrated with EPON Araldite (Electron Microscopy Sciences) and polymerized overnight at 60 C. Serial semi-thick sections (200 nm) were cut using an Ultracut UCT Microtome (Leica Microsystems, Vienna, Austria). Sections were collected on piliform-coated copper slot grids and poststained with 2% uranyl acetate in 70% methanol, followed by Reynold’s lead citrate57.

### Data acquisition by electron tomography

Colloidal gold particles (15 nm; Sigma-Aldrich) were attached to both sides of semi-thick sections collected on copper slot grids to serve as fiducial markers for subsequent image alignment. For dual-axis electron tomography, a series of tilted views were recorded using a TECNAI F20 transmission electron microscope (FEI Company, Eindhoven, The Netherlands) operated at 200 kV. Images (4K x 4K) were captured every 1h over a ±60 range and a pixel size of 2.3nm using a TVIPS CCD camera. We used the IMOD software package (http://bio3d.colourado.edu/imod) for image processing, which contains all the programs needed to calculate electron tomograms. The tilted views were aligned using the positions of the colloidal gold particles as fiducial markers. Tomograms were computed for each tilt axis using the R-weighted back-projection algorithm. Reconstructed tomograms were flattened.

### Image analysis, quantification, and statistics

All the experiments were repeated three times. All the measurements and analyses were performed on at least three embryos. Sample size, indicated by “n” values, is included in the figure legends. For length measurement (cilia length, ciliary tip length, and TZ length), at least three cells per embryo were chosen randomly; however, it was ensured that only those cilia/signals that were distinct and not overlapping (or tangled) with neighboring ones were measured. For statistical analysis, Prism ver. 10 was used. The type of analysis, p-values, and significance are included in the figure legends. The F-actin intensity measurements were done using Fiji. The cortical actin refers to the actin network along the MCC boundary, while the medial actin refers to the F-actin network at the apical surface of the MCC. The intensity measurements were performed using a line or polygon tool in Fiji, as described in an earlier study^20^. For orientation calculations, images were taken after aligning the embryos horizontally from anterior on the left to posterior on the right and stained for Clamp-GFP (rootlet) and Centrin-RFP (basal body). The Clamp-GFP orientations were calculated in MATLAB R2021a using image processing and circular statistics toolboxes. The cilia orientations were drawn by hand from rootlet to basal body using the ‘drawline’ command. The angles were calculated with respect to the global X-axis in the cartesian coordinate system (first point starting at 0,0) using the command ‘atan2d’, which gives the angles from 0° to +180° in a counterclockwise direction (quadrants I and II) and 0° to -180° in a clockwise direction (Quadrants IV and III). 26 and 21 multiciliated cells were analyzed for control and deciliated embryos, respectively, with an average of 112 cilia orientations (∼60-200 cilia per cell) in each cell for each condition. For the IFT-GFP (IFT20, IFT80, and IFT43) fluorescence intensity calculations, we adopted an unbiased approach to measure the IFT intensities. Firstly, the three channels were separated using Fiji. The cilia and centrin channels were merged, and a circular ROI between 0.646-0.680 μm^2^ was drawn around the ciliated and non-ciliated centrin-RFP puncta (4-5 basal bodies from each category). The mean gray value of the IFT-GFP channel was calculated, and the values were normalized from 0-1. The normalization was done using the following formula:

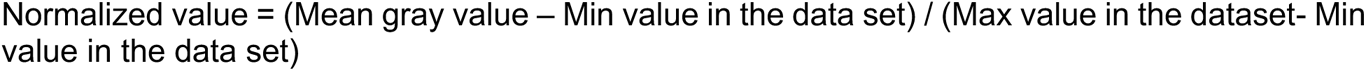

### Theoretical estimate of force applied by a single Multiciliated Cell (MCC)

Boselli et al. derived a simple force model for a cilium from resistive force theory ^58^using a simplified assumption of cilium as a rigid rod rotating around the base, making an arc of varying lengths from 0 to length of the cilium (l) during each cycle. By ignoring force contribution during the recovery stroke, the effective force contribution from a cilium during a cycle is simplified as

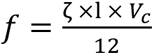

where l is the length of the cilium,

V_c_ is the velocity approximated as ∼4×ν×l where ν is the beat frequency (assumed as 25 Hz and l is the length of the cilium) and

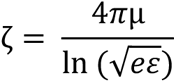

is the transverse drag coefficient,

where µ is the viscosity of the liquid (assumed as 0.001 Pa. s for water)

e is the natural log (e=2.718281828459)

*ε* is the cilium aspect ratio (length/thickness), the thickness of the cilium is assumed as 0.2 µm.

Force contribution from the entire MCC is approximated as F = f × N × C_o_

where f is the force contribution from a single cilium where N is the total number of cilia in the cell, and C_o_ is the coupling coefficient, which is defined as 0.55 based on

Boselli et al.’s force calculations^58^ for far-field estimates show that ∼50-60 % of cilia don’t contribute to a cell’s final average lateral force component due to entanglement and phase shifts^58^. Hence, a value of C_o_ = 0.55 is used for the calculations, reflecting close to the actual force contribution from each cell. Using the formula described force contribution from a single MCC was simulated for varying lengths (1-15 µm assuming all cilia in the cell have the same length) and number of cilia (1-150) and visualized as surface plots.

## Key resource table

| **Antibodies** |  |  |  |
| --- | --- | --- | --- |
| **Name** | **Marker** | **Make and Cat. No.** | **Remark** |
| Mouse Anti-Acetylated α-tubulin (6-11B1) | Ciliary axoneme | Sigma | PFA fix, 1:1000 |
| Rabbit mAb Acetyl-α-Tubulin (Lys40) (D20G3) | Ciliary axoneme | Cell Signaling technology (#5335) | Methanol Fix, 1: 500 |
| Rabbit mAb Acetyl-α-Tubulin (Lys40) (D20G3) (Alexa Fluor® 647 Conjugate) | Ciliary axoneme | Cell Signaling Technology (#81502) | PFA fix, 1: 25 |
| Mouse pAb Anti-B9d1 | Transition Zone | Sigma (SAB1409114) | Methanol Fix, 1:50 |
| Chicken anti Mouse conjugated to Alexa fluor 488 | NA | Invitrogen | 1:500 |
| Anti-Rabbit conjugated to Alexa fluor 647 | NA | Jackson | 1:500 |
| Phalloidin conjugated to Alexa fluor 488 | Actin | Invitrogen | 1:50 |
| Phalloidin conjugated to Alexa fluor 647 | Actin | Invitrogen | 1:50 |
| **DNA/RNA** |  |  |  |
| **Name** | **Marker** | **Conc. injected** | **Source** |
| Chibby-tagBFP2 | Basal body | 100 pg (RNA) | This study |
| mScarlet-Sentan | Cilia tip | 200 pg(RNA) | This study |
| GFP-Clamp | Rootlets | 200 pg (RNA) | <sup>69</sup> |
| Centrin-RFP | Basal body | 100 pg (RNA) | <sup>69</sup> |
| MemRFP | Cell and Ciliary membrane | 100 pg(RNA) | <sup>50</sup> |
| IFT43-GFP | Cilia base/Cilia | 150pg (RNA) | <sup>27</sup> |
| IFT20-GFP | Cilia base/Cilia | 150pg (RNA) | <sup>27</sup> |
| IFT80-GFP | Cilia base/Cilia | 150pg (RNA) | <sup>27</sup> |
| **Drugs/Chemicals** |  |  |  |
| **Name** | **Solvent** | **Make and Cat No.** | **Remark** |
| Cycloheximide | DMSO | Sigma | 10µg/ml |
| MG132 | DMSO | Sigma | 10 µM |

## ACKNOWLEDGMENTS

The authors would like to express their gratitude to Dr. Bob Bloodgood and the members of the Kulkarni Lab for their invaluable feedback and thorough review of this manuscript. Appreciation is also extended to Dr. Mustafa Khokha for the initial discussions and intellectual guidance for the project. Acknowledgment is also due to the Molecular Electron Microscopy Core and the Advanced Microscopy Facility at the University of Virginia. Lastly, we are grateful for the NIH Pathway to Independence, K99/R00 (K99HL133606 and R00HL133606) grant awarded to Saurabh Kulkarni.

## SUPPLEMENTARY FIGURE LEGENDS

**Figure S1:**
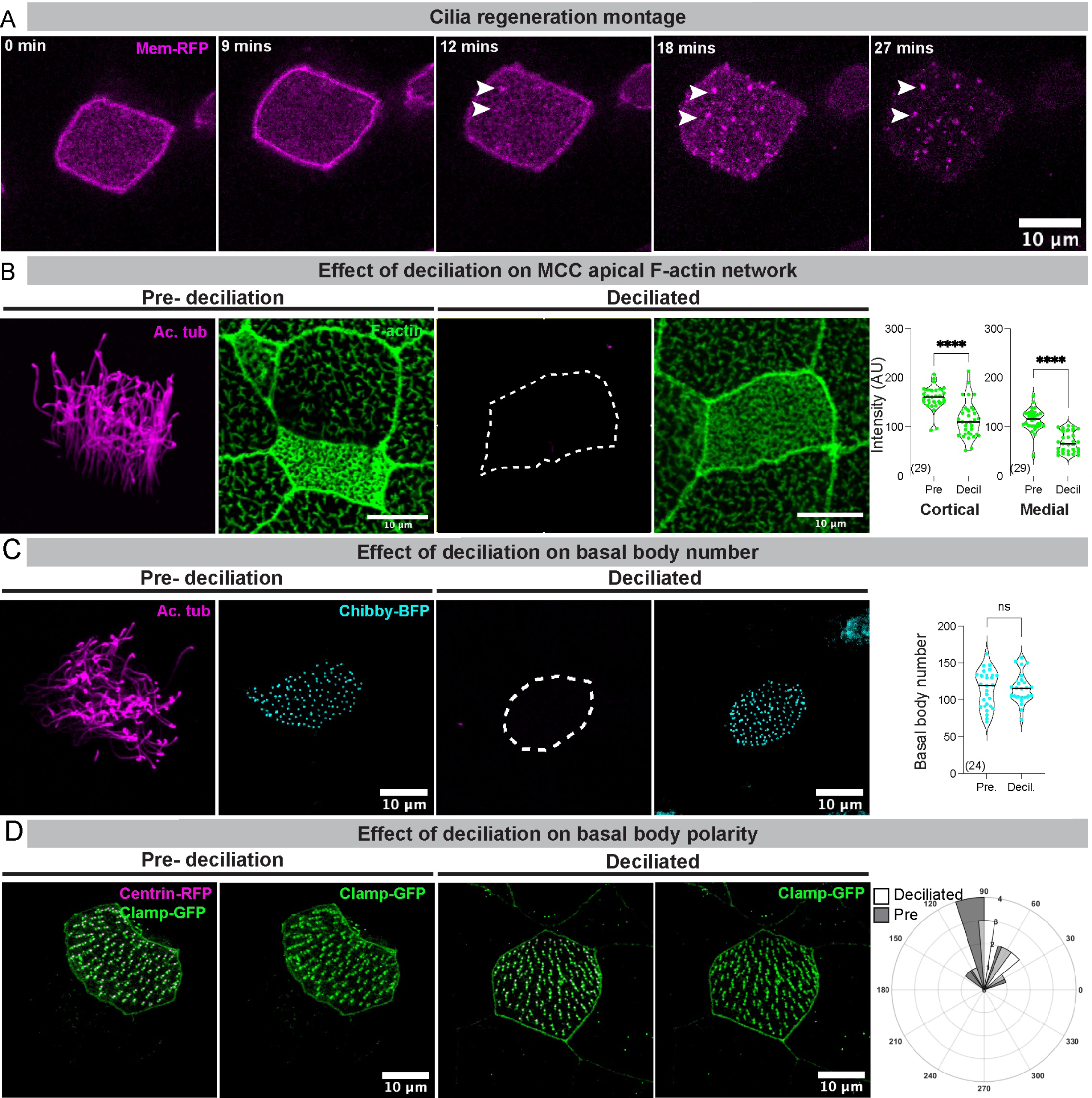
Deciliation affects apical F-actin, but the basal body number is unaffected. A) Montage of regenerating cilia (mem-RFP) in the animal caps. The mem-RFP signal specks can be seen emerging from the cell’s surface by ∼12 mins, eventually growing into beating cilia (see Supplement video 1). B) Pre- and post-deciliation MCCs that are stained for cilia(magenta) and F-actin(green) are depicted. Cortical and medial F-actin intensity significantly differs in Pre and 0-hr. deciliated samples. The values in parenthesis indicate the number of MCCs measured from three trials from 9-10 embryos. **** - p<0.0001, Mann-Whitney test. C) The number of basal bodies labeled with Chibby-GFP (cyan) is unaffected after deciliation. The values in parenthesis indicate the number of MCCs measured from three trials from 9-10 embryos. ns = not significant, Mann-Whitney test. D) Deciliation does not affect basal body polarity. Basal body polarity was determined by measuring the orientation of rootlets labeled with Clamp-GFP (green) with their respective basal body labeled with Centrin-RFP (magenta) and represented in the rose plot. The directionality was measured in 27 pre-deciliated cells and 22 MCCs post-deciliation from three trials from 9-10 embryos in each category.

**Figure S2:**
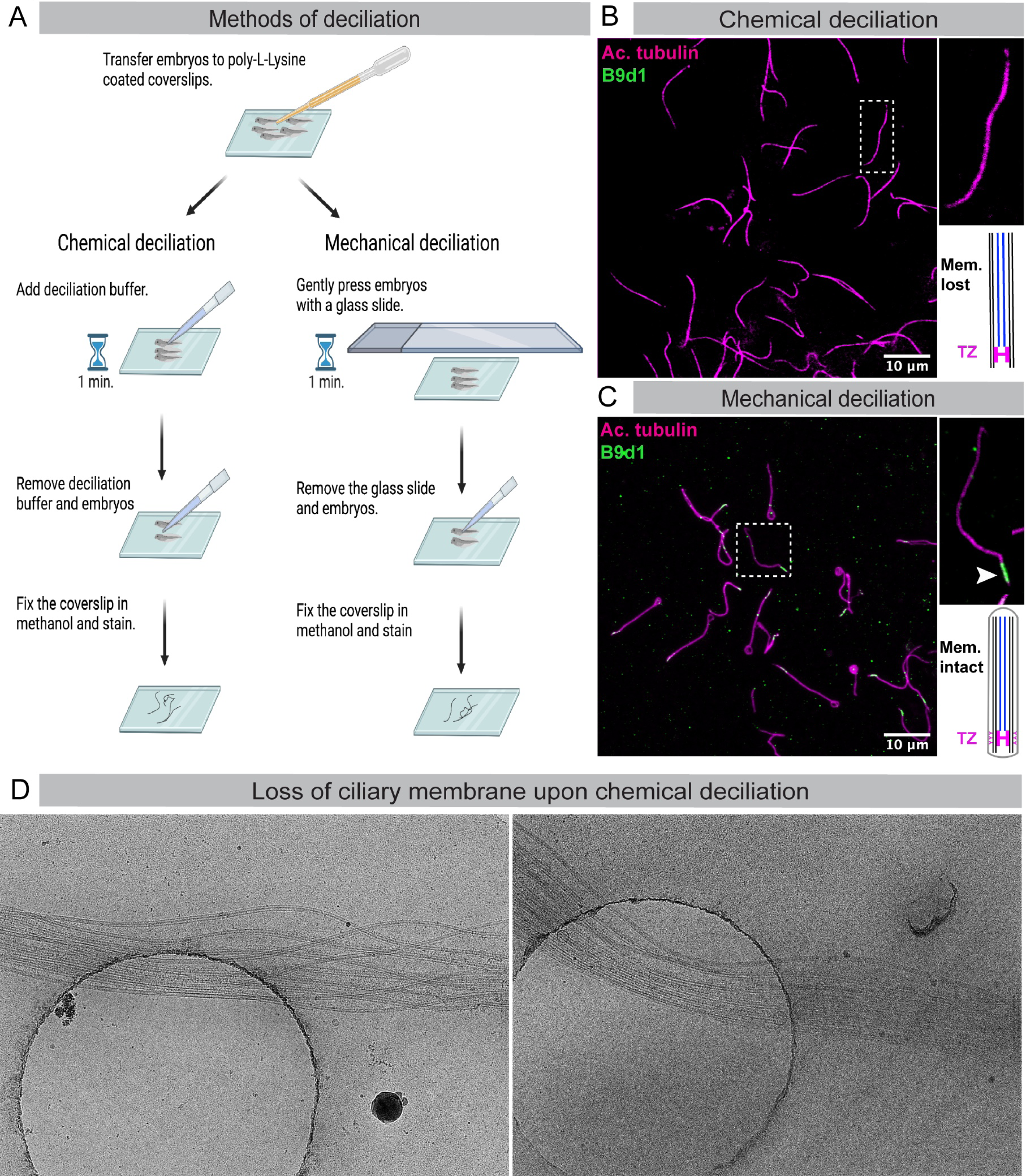
Transition zone is removed with cilia during deciliation. A) Schematic of the chemical and mechanical deciliation methods. B) Cilia with chemical deciliation lose the B9d1 signal, possibly due to the loss of membrane with detergent in the deciliation buffer, whereas C) the B9d1 signal is maintained with mechanical deciliation. D) Electron micrographs of cilia from the chemical deciliation method lack the ciliary membrane and show splaying of axonemal microtubules.

**Figure S3:**
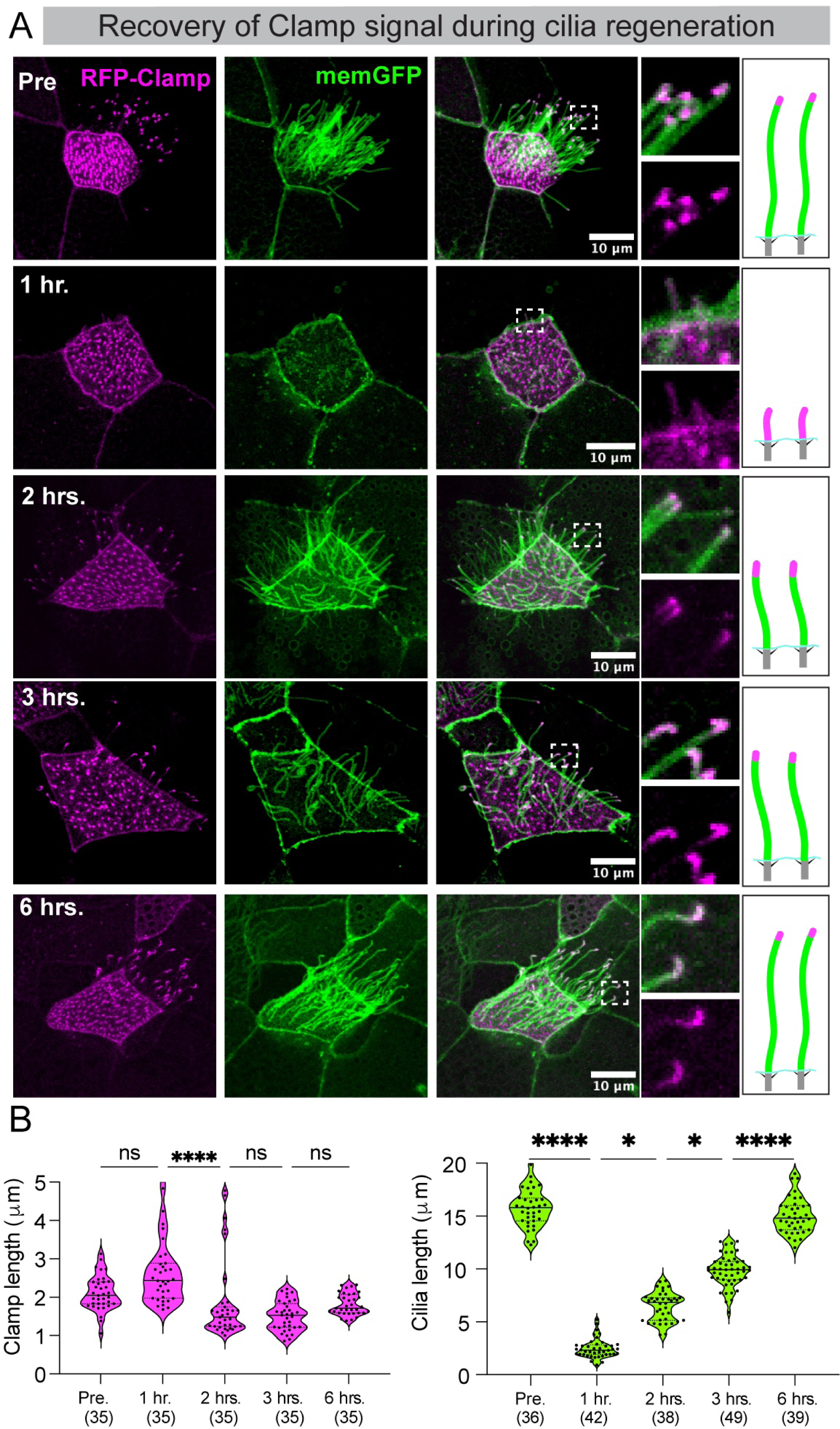
Clamp is localized to the ciliary axoneme during the early stages of regeneration. B) MCCs are labeled with RFP-Clamp (ciliary tip and base, magenta) and memGFP (cilia, green) at various stages of cilia regeneration. After 1 hr. post-deciliation, Clamp signal can be seen in the ciliary axonemes. At 2 hrs., Clamp signal starts accumulating at the ciliary tips. At 3 and 6 hr., Clamp signal appears more like pre-deciliated samples. B) Clamp signal length (left panel) and cilia length (right panel) were measured and compared among different time points. The values in parenthesis indicate the number of cilia measured from three trials using 9-10 embryos. **** - p<0.0001; * - p< 0.05; ns - not significant, Kruskal-Wallis test, followed by Dunn’s test.

**Figure S4:**
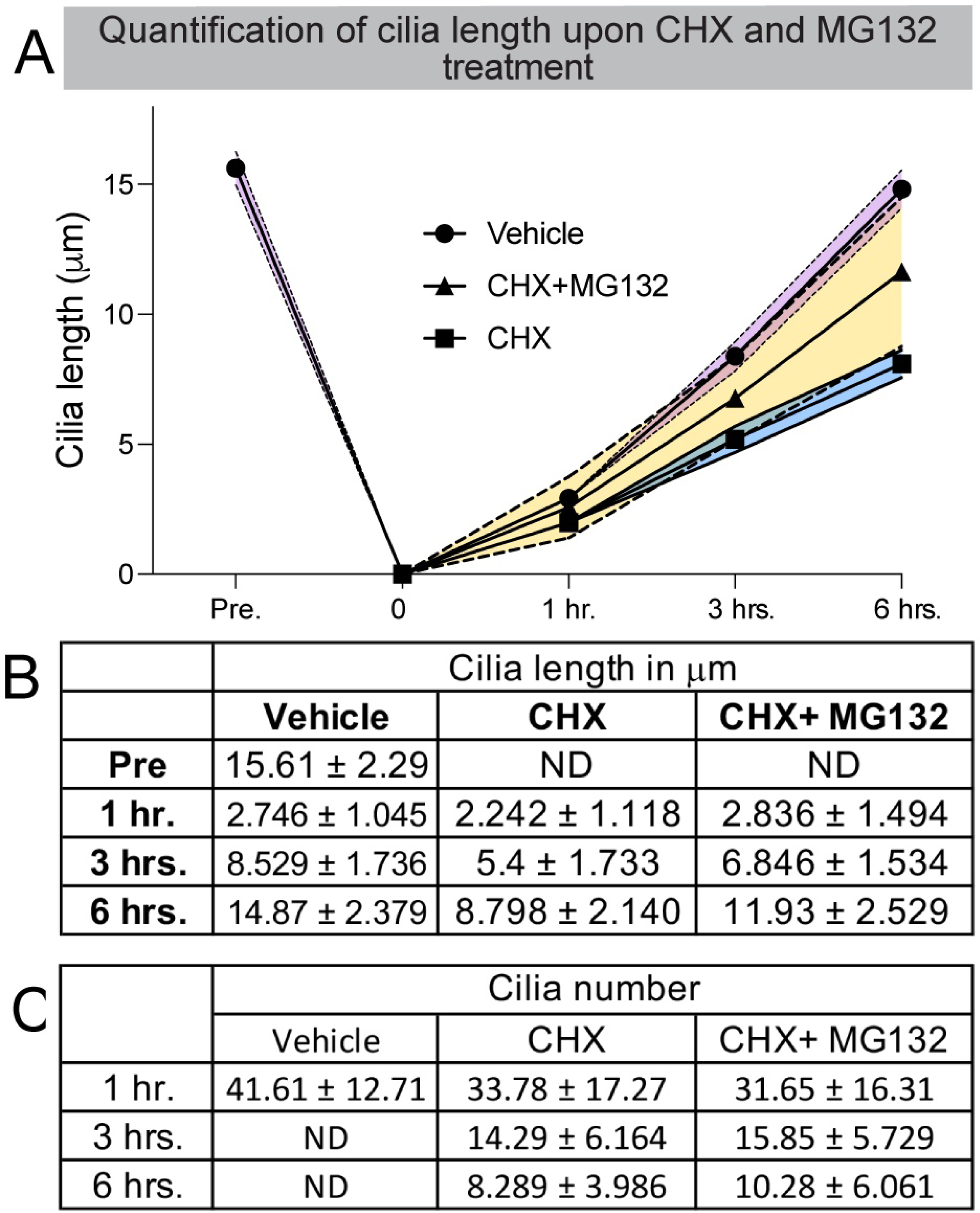
Quantification of cilia length and number on CHX and CHX+MG132 treatment. A) Graph depicting the length of regenerating cilia in the vehicle, CHX, and CHX+MG132 treatments. The cilia lengths at every time point, treatment, and the ‘n’ values are shown in Fig 5D. B) The Table shows the average cilia length in 3 treatments and 4 time points. Values are represented as mean±SD. The “n” values are shown in Fig. 5D. C) Table showing variation in cilia number in 3 treatments and 3 time points. Values are represented as mean±SD. The n values and statistical differences are shown in Fig. 5E. ND— Not determined.

**Figure S5:**
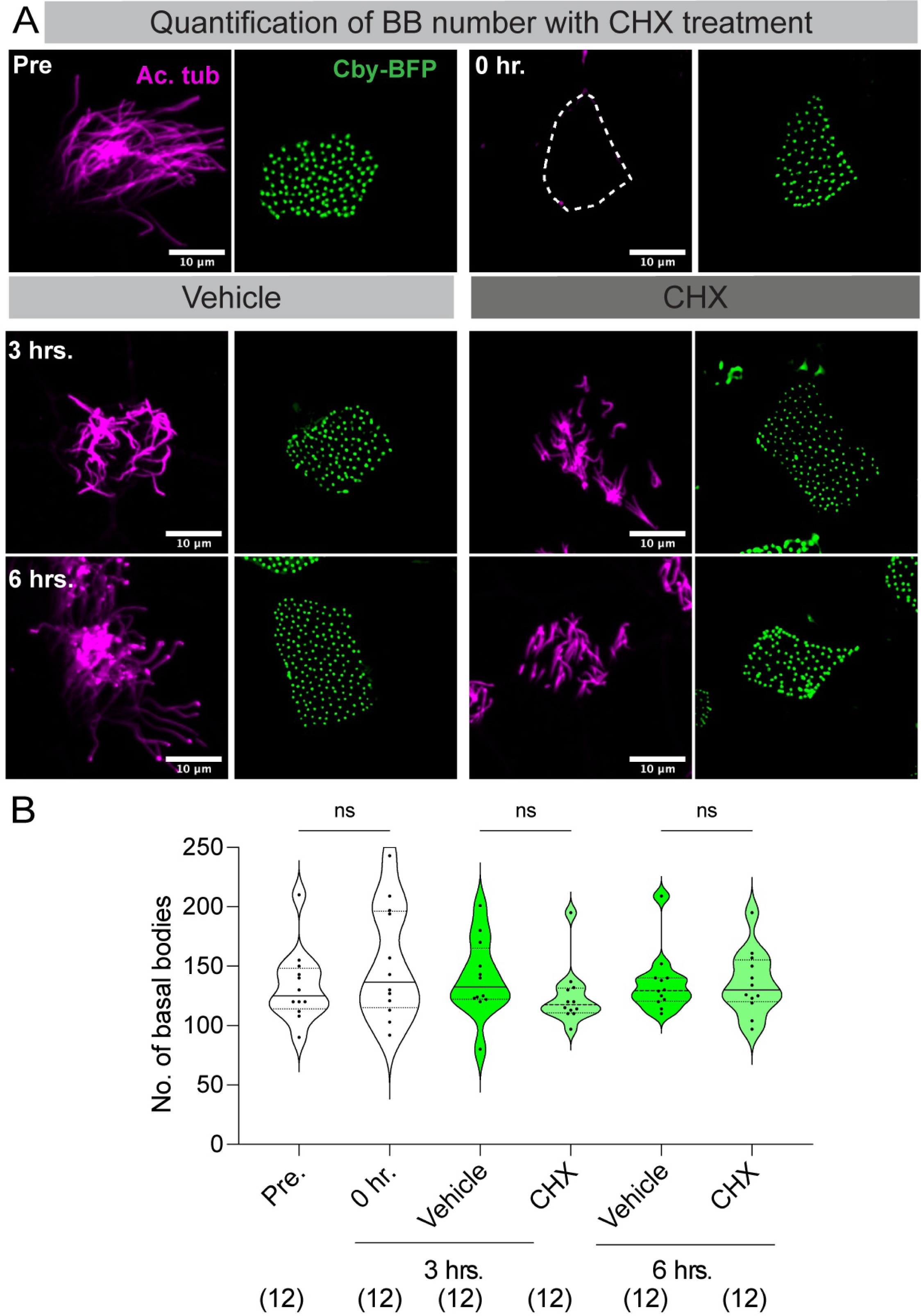
Blocking new protein synthesis does not affect the basal body during regeneration. A) Stage 28 embryos labeled Chibby-BFP (green, basal bodies) were transferred to DMSO containing 1/9 MR (vehicle) or cycloheximide (CHX) containing 1/9 MR post-deciliation. The embryos were allowed to regenerate and collected at 3 and 6 hrs. Embryos were fixed and stained for cilia (magenta) using Ac. α-Tubulin. B) The Chibby puncta were counted in all samples (deciliation and regeneration), and the quantification of the basal body number is shown in the graph. The values in parenthesis indicate the number of cells counted from 8 embryos from three trials. ns - not significant, Kruskal-Wallis test, followed by Dunn’s test.

**Figure S6:**
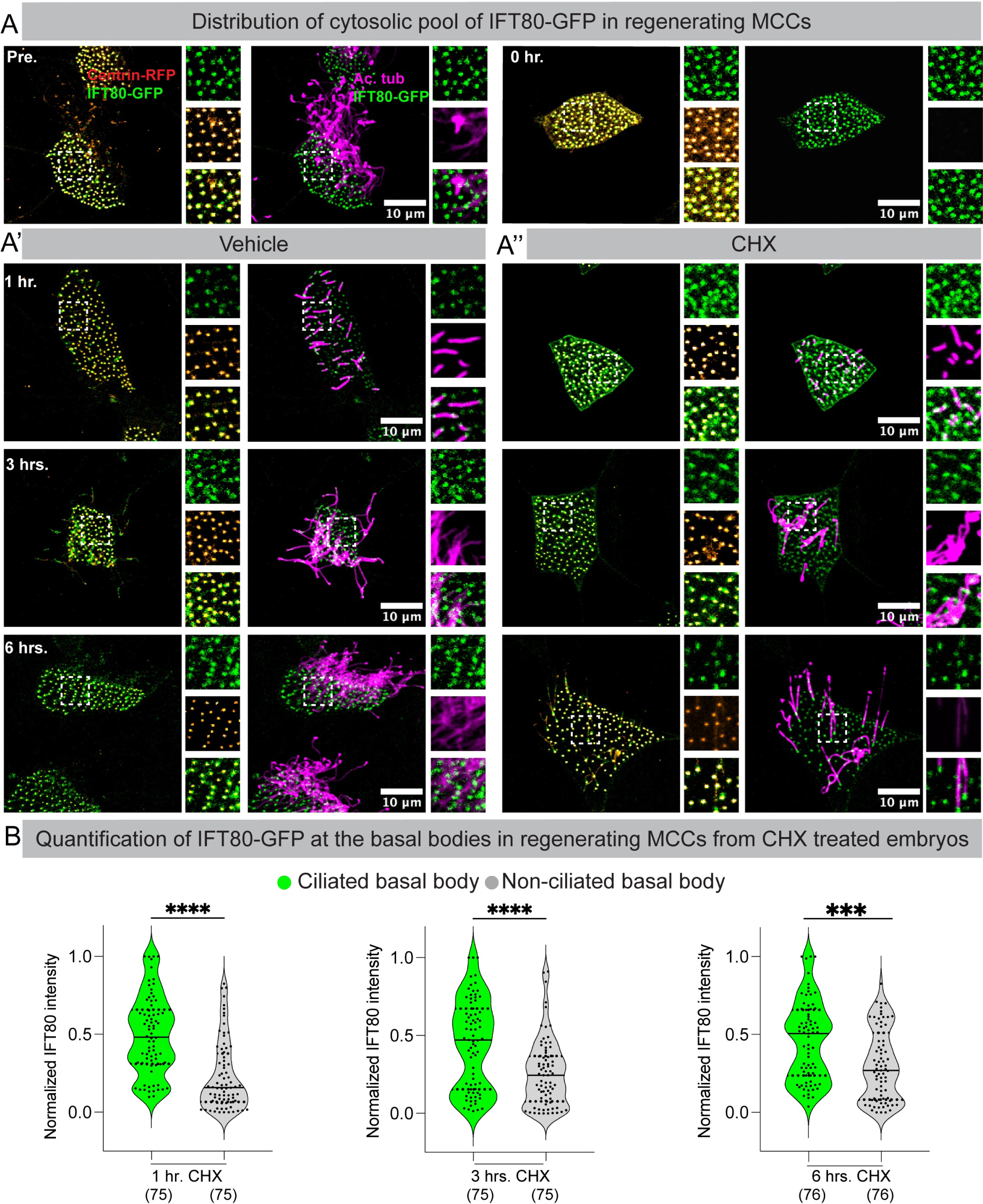
Distribution of ciliary precursor pool (IFT80-GFP) in MCCs. A) Embryos injected with IFT80-GFP (green) and Centrin-RFP (orange hot, basal bodies) were deciliated at stage 28 (0 hr.) and were split into two experiments (DMSO and CHX). A’, A’’) The embryos in both sets regenerated cilia for 6 hours. After fixation, the embryos were stained for cilia(magenta). Note that the control MCCs at all time points (1 hr., 3 hrs., and 6 hrs.) have multiple cilia regenerating, and the IFT80-GFP intensity is uniform at every basal body. In contrast, the number of regenerating cilia decreases with time in CHX-treated samples, and the IFT80-GFP is enriched at a few ciliated basal bodies. B) The intensity of IFT80-GFP associated with ciliated (green) vs. non-ciliated(gray) basal bodies in the same MCC (in CHX-treated samples) at different time points during cilia regeneration. A total of 8-10 basal bodies per MCC (4-5 ciliated and 4-5 non-ciliated) and 5 MCCs were chosen, and the mean gray value was estimated and normalized to the maximum and the minimum values in the set. The value in parenthesis indicates the number of basal bodies analyzed (with and without IFT80-GFP) from 9 embryos from three independent trials. Note the significant difference in the IFT80-GFP signal intensity at ciliated vs. non-ciliated basal bodies at all time points. **** - p<0.0001, Kruskal Wallis test, followed by Dunn’s multiple comparison test.

**Figure S7:**
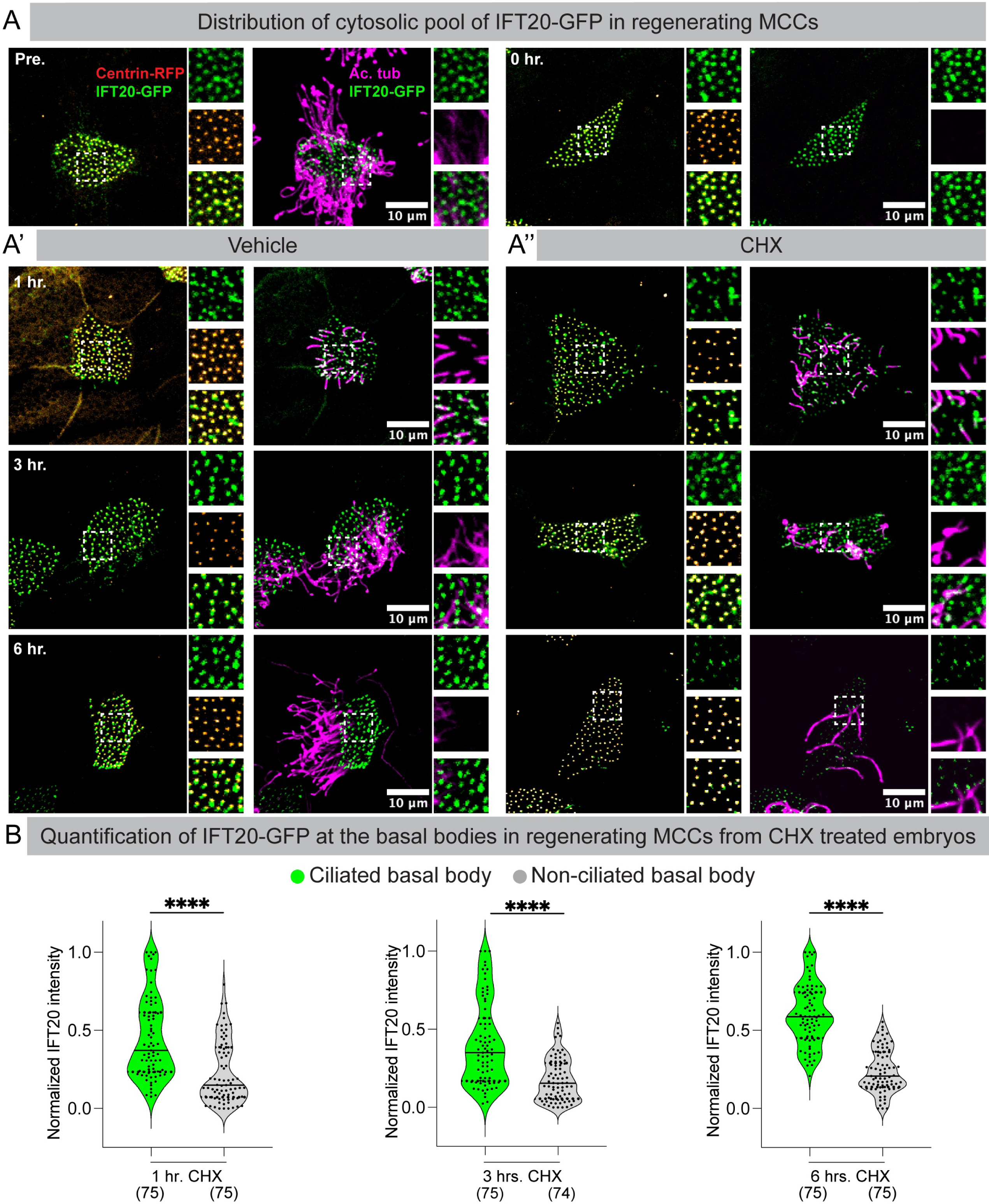
Distribution of ciliary precursor pool (IFT20-GFP) in MCCs. A) Embryos injected with IFT20-GFP (green) and Centrin-RFP (orange hot, basal bodies) were deciliated at stage 28 (0 hr.) and were split into two experiments (DMSO and CHX). A’, A’’) The embryos in both sets regenerated cilia for 6 hours. After fixation, the embryos were stained for cilia(magenta). Note that the control MCCs at all time points (1 hr., 3 hrs., and 6 hrs.) have multiple cilia regenerating, and the IFT20-GFP intensity is uniform at every basal body. In contrast, the number of regenerating cilia decreases with time in CHX-treated samples, and the IFT20-GFP is enriched at a few ciliated basal bodies. B) The intensity of IFT20-GFP associated with ciliated (green) vs. non-ciliated(gray) basal bodies in the same MCC (in CHX-treated samples) at different time points during cilia regeneration. A total of 8-10 basal bodies per MCC (4-5 ciliated and 4-5 non-ciliated) and 5 MCCs were chosen, and the mean gray value was estimated and normalized to the maximum and the minimum values in the set. The value in parenthesis indicates the number of basal bodies analyzed (with and without IFT20-GFP) from 9 embryos from three independent trials. Note the significant difference in the IFT20-GFP signal intensity at ciliated vs. non-ciliated basal bodies at all time points. **** - p<0.0001, Kruskal Wallis test, followed by Dunn’s multiple comparison test.

## SUPPLEMENTARY VIDEO LEGENDS

**Supplement Video 1: Mucociliary epithelium regenerates cilia in the same MCC.**

Live imaging of animal caps dissected from embryos injected with membrane-RFP RNA. Imaging was started approximately 2 minutes post-deciliation. The time stamp on the video is when the imaging was started and not the time of deciliation.

**Supplement video 2 - 6: Cilia regeneration tomogram data.**

Tomograms from control and regenerating MCCs. Cilium from the control sample shows the presence of TZ, indicated by an ‘H’ shaped electron-dense structure (video 2). Note that this structure is missing in samples after 20 minutes of cilia regeneration (video 3) and begins to appear after 1 hour (video 4). By the end of 3 hours (video 5) and 6 hours (video 6) of cilia regeneration, the tomograms reveal complete regeneration of the transition zone.

## SUPPLEMENTARY DATA TABLE

**Table 1:** Force data computed for different cilia numbers and lengths with maximum length of 15um and maximum number of 150 cilia.

